# A more complex basal complex: novel components mapped to the *Toxoplasma gondii* cytokinesis machinery portray an expanded hierarchy of its assembly and function

**DOI:** 10.1101/2021.10.14.464364

**Authors:** Klemens Engelberg, Tyler Bechtel, Cynthia Michaud, Eranthie Weerapana, Marc-Jan Gubbels

## Abstract

The basal complex (BC) of *Toxoplasma gondii* has an essential role in cell division but details on the mechanism are lacking. To promote insights in this process, reciprocal proximity based biotinylation was used to map the basal complex proteome. An assembled protein map was interrogated by spatiotemporal characterization of critical components as well as functionally by disrupting the expression of the components. Spatially, this revealed four proteins sub-complexes with distinct sub-structural BC localization. Temporally, several patterns were differentiated based on their first appearance and/or disappearance from the BC corresponding with different steps in BC development (initiation, expansion, constriction, maturation). We also identified a protein pre-ceding BC formation (BCC0) laid out in a 5-fold symmetry. This symmetry marks the apical annuli and site of alveolar suture formation. From here, it was determined that the apical cap is assembled in the apical direction, whereas the rest of the IMC expands in the basal direction, inspiring a new bi-directional daughter budding process. Furthermore, we discovered BCC4, an essential protein exclusively localizing to the BC during cell division. Although depletion of BCC4 did not prevent BC formation, it led to BC fragmentation at the mid-point of cell division. Based on these data, a model is presented wherein BCC4 and MORN1 stabilize each other and form a rubber band that implies an essential role for the BC in preventing the fraying of the basal end of the assembling daughter cytoskeleton scaffolds. Furthermore, one new component of the Myosin J and Centrin2 cluster was BCC1, a hypothetical protein whose depletion prevents the non-essential last step of BC constriction. Overall, the BC is a highly dynamic, multi-functional structure that is critical to the hierarchical assembly of the daughter parasites.

## Introduction

The Apicomplexa are obligate intracellular parasites infecting a wide range of hosts, including several species that infect humans and cause significant diseases. *Toxoplasma gondii* is widespread and has infected one third of the global human population. Although most infections are chronically dormant without disease symptoms, opportunistic infections lead to a spectrum of clinical manifestations in the immunocompromised, immunosuppressed or individuals with an immature immune system (Montoya and Liesenfeld, 2004). All pathology originates from tissue destructions caused by fast rounds of lytic, intracellular cell divisions of the acute, tachyzoite life stage.

*T. gondii* cell division differs in many respects from the conventions of mammalian cell division. Tachyzoites divide by budding two daughter cells inside the mother cell (i.e. endodyogeny = internal budding). In this process daughter cytoskeleton scaffolds nucleate on the duplicated centrosomes and grow in an apical-to-basal direction (Anderson-White et al., 2012; Chen and Gubbels, 2013; Francia and Striepen, 2014; Suvorova et al., 2015). Many of the cytoskeleton components are unique to the parasite and absent from the mammalian host, such as a family of intermediate filament-like, alveolar domain containing proteins associated with alveolar vesicles making up the inner membrane complex (IMC) (Anderson-White et al., 2011; Anderson-White et al., 2012). The cortical membrane skeleton is principally different from the actin-spectrin cytoskeleton in mammals and imposes distinct needs on the secretory system as well as the cell division machinery (Goodenough et al., 2018). The membrane skeleton is buttressed by a set of 22 subpellicular microtubules emanating from the apical end, which itself is capped by a unique microtubular basket known as the conoid.

Cell division starts with Golgi duplication, followed by centrosome duplication and assembly of cytoskeleton elements on outer core of the duplicated centrosomes (Nishi et al., 2008; Suvorova et al., 2015). The early formation of daughter cytoskeletal scaffolds is marked by simultaneous recruitment of an F-BOX ubiquitin ligase complex protein (FBXO1) (Baptista et al., 2019) and two unique apicomplexan proteins, apical cap protein 9 (AC9) (O’Shaughnessy et al., 2020; Tosetti et al., 2020) and IMC32 (Torres et al., 2021) which lay down the basic connections between the three major cytoskeleton components: microtubules, alveolar vesicles, alveolin-domain IMC proteins. Subsequently, the conoid, an apical microtubular basket, is formed in the center of this ring (Tosetti et al., 2020), and from here on the membrane cytoskeleton grows in an apical to basal direction by adding components on the basal end (Anderson-White et al., 2012).

A ring structure situated on the very basal, posterior edge of the daughter scaffolds is known as the basal complex (BC) and is essential to complete cell division (Gubbels et al., 2006; Hu et al., 2006; Heaslip et al., 2010; Lorestani et al., 2010). The BC functions as the contractile ring during cell division (Gubbels et al., 2006; Lorestani et al., 2010). MORN1 was identified as key scaffolding component of the BC (Lorestani et al., 2010; Lorestani et al., 2012). Surprisingly, *T. gondii* completes cell division in presence of actin (Shaw et al., 2000; Periz et al., 2017), unlike the actin-dependent contractile ring of the mammalian host. However, it has been established that Myosin J (MyoJ) is responsible for the last step of BC constriction, that this step actually is actin dependent, but that parasites lacking MyoJ only display mild fitness loss (Frenal et al., 2017). Yet, interfering with BC assembly at earlier steps induces lethal phenotypes, either by overexpressing the BC scaffolding protein MORN1 (Gubbels et al., 2006), or MORN1 depletion resulting in daughter scaffolds that are much wider open and result in conjoined, double- or multi-headed parasites (Lorestani et al., 2010). Thus, the early assembly of the BC is essential to complete cell division, which seems to imply that there is another actin-myosin-independent process acting early in cell division.

The inventory of proteins that thus far have been mapped to the BC does not immediately highlight the unconventional mechanism of its essential function during cell division. Nevertheless, we have provided detailed insights into the architecture and spatio-temporal dynamics of the BC. Four groups of spatiotemporally defined IMC proteins are sequentially recruited and serve as a set of highly-resolved daughter development markers (Anderson-White et al., 2011; Anderson-White et al., 2012). Scaffolding protein MORN1 is deposited already at formation of the daughter bud (Gubbels et al., 2006). Halfway through daughter development additional proteins are recruited to the BC such as the intermediate filament-like alveolins IMC5, IMC8, IMC9 and IMC13 (Anderson-White et al., 2011) as well as MyoJ (Frenal et al., 2017) and Centrin2 (Cen2) (Hu et al., 2006; Hu, 2008). This recruitment coincides with the onset of tapering the cytoskeleton scaffolds to the posterior end (Hu, 2008; Anderson-White et al., 2011). A last transition in BC composition occurs upon daughter parasite emergence when proteins such as FIKK (Skariah et al., 2016) and MSC1a (Lorestani et al., 2012) are recruited. The role of these proteins, which are only found in the BC of the mother cell, is currently unknown. The set of known BC markers in combination with electron microscopy resolves into three substructures within the BC based on a combination of molecular and ultrastructural data (Mann and Beckers, 2001; Hu, 2008; Anderson-White et al., 2011) (Fig. 1a).

**Fig 1.**
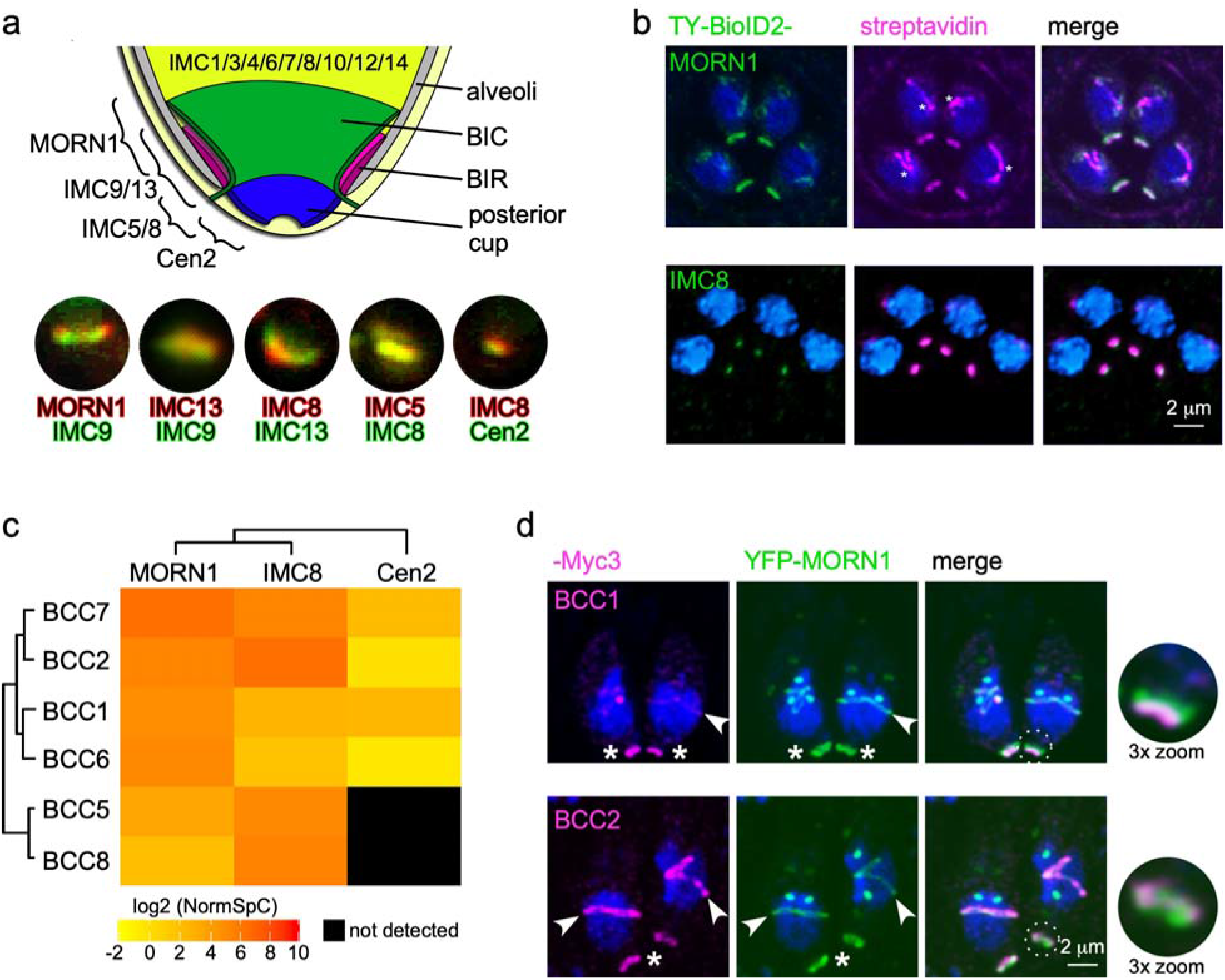
Proximity biotinylation reveals novel BC components. **a.** Schematic of the *T. gondii* BC highlighting distinct localizations of the utilized BioID-baits. Double-stained panels from (Anderson-White et al., 2011). **b.** Endogenously tagged BioID2 parasite lines in the MORN1 or IMC8 locus, show correct localization and biotinylation capability in presence of 150 μM biotin for 16 hrs. **c.** Abundance of novel basal complex components (BCCs) components across BioID2 experiments with the indicated, known BC components as baits. The heat map depicts normalized spectral counts (NormSpC) for each BCC prey. BCCs were numbered in order of their validation. **d.** Examples of subcellular localization analysis of endogenously C-terminal 3xMyc-tagged BCCs co-stained with YFP-MORN1 to confirm BC localization. Distinct temporal localization kinetics to the mother (asterisks) and daughter BC (arrow heads) are revealed, as well as distinct spatial patterns relative to MORN1 in the mature BC (3xzoom panels of the regions marked with the dotted lines).

Finally, it is of note that both MORN1 and Cen2 have additional localizations in the cell: MORN1 in the spindle pole and apical end of the cytoskeleton (Gubbels et al., 2006) and Cen2 in the centrosome, apical annuli and apical polar ring (Hu et al., 2006). Overall, the parts list lacks leads toward a putative restrictive mechanism in the BC prior to the final constriction by MyoJ/Cen2.

Toward a mechanism of the essential function of the BC and to expand our knowledge on *T. gondii* cell division we applied reciprocal BioID using six different baits. Statistical analysis and experimental validation identified 10 novel BC components (BCC1-10) resolving across four BC sub clusters (BCSC1-4) correlating with the previously identified architecture (Anderson-White et al., 2011). In addition, we identified a protein preceding BC formation (BCC0) that laid out a 5-fold symmetry conserved throughout the cytoskeleton in the apical annuli and alveolar vesicle architecture. Moreover, our data showed that the daughter buds bi-directional from this structure. In addition, we discovered that BCC4 depletion phenocopies MORN1 depletion, is not essential for BC formation, but is required to maintain the structural integrity of nascent daughter buds. This led us to propose a ‘rubber band’ model that mechanistically explains the essential role of the BC in early stages of cell division. Overall, our work uncovered a novel dimension to the hierarchy and structure of daughter budding as well as answered the long-standing question of the essential function of the BC during cell division.

## Results

### Mapping the basal complex by reciprocal proximity biotinylation

We previously defined distinct compartments in the BC (Anderson-White et al., 2011), and used three baits representing these compartments as starting point for mapping the BC proteome by proximity biotinylation (Fig. 1a). MORN1, IMC8 and Centrin2 (Cen2) represent the widest upper part, the middle section and the lower section of the BC, respectively, while noting that there is partial overlap between these localization patterns (Hu, 2008; Anderson-White et al., 2011). We fused the endogenous ORFs of these genes at their 5’-end with a Ty-tagged BioID2 and confirmed the correct localization and biotinylation capacity by fluorescence microscopy (Fig. 1b; Ty-BioID2-Cen2 was reported before (Engelberg et al., 2020)). Following mass spectrometry, we mined the data for prey abundance by calculating length-normalized spectral counts (NormSpC) (Youn et al., 2018; Go et al., 2021). We screened for novel BCCs by ranking NormSpC values from the highest to the lowest in each experiment and focused on the top 25 hits. Besides known BC proteins and hits with known other localizations, this list contained six hypothetical proteins which we named putative BC components (BCC) followed by a number representing the order in which we validated them: TGGT1_232780 (BCC1), TGGT1_231070 (BCC2), TGGT1_269460 (BCC5), TGGT1_202550 (BCC6), TGGT1_311230 (BCC7), TGGT1_273050 (BCC8) (Fig. 1c, Table 1). Different NormSpC values were seen for these candidates for bait, potentially reflecting the physical distance between bait and prey in the BC. BCC5 and BCC8 were not present in the Cen2 data set. BC localization for these six new BCCs was validated by tagging the endogenous genes with a triple Myc or YFP-tag, which also highlighted distinct spatiotemporal localization patterns (Fig. 1d, Suppl. Fig. S1). Taken together, proximity-labeling successfully recovered known BC proteins as well as identified a set of novel BCC proteins associated with the BC.

**Table 1.** All proteins mapped to the BC in this study. BCSC assignments based on averaged statistical analysis combined with experimental validation. Data derived from ToxoDB (Gajria et al., 2008), which harbors the primary data reported as follows: phosphoproteome (Treeck et al., 2011), lytic cycle fitness score (Sidik et al., 2016), hyperLOPIT subcellular localization assignments (Barylyuk et al., 2020). * Nuclear LIM Interacting factor family phosphatase.

| BC-SC# | name | TGGT1 | annotation / domains | fitness | MW (kDa) | # aa | #P-sites | # exons | LOPIT |  |
| --- | --- | --- | --- | --- | --- | --- | --- | --- | --- | --- |
|  |  |  |  |  |  |  |  |  | MAP | MCMC |
| (1) | BCC0 | 294860 | hypothetical, N-term acetyl. | -4.10 | 255 | 2457 | 14 | 5 | N/A | N/A |
| 4 | BCC1 | 232780 | hypothetical | -1.88 | 78 | 713 | 7 | 8 | DG | PM-p2 |
| 2 | BCC2 | 231070 | protein kinase | 0.22 | 141 | 1318 | 40 | 1 | IMC | IMC |
| 2 | BCC3 | 311770 | hypothetical | -0.11 | 223 | 2076 | 57 | 2 | N/A | N/A |
| 1 | BCC4 | 229260 | hypothetical | -3.52 | 364 | 3460 | 46 | 1 | N/A | N/A |
| 3 | BCC5 | 269460 | S/T phosphatase; EF-hand | -0.81 | 219 | 2066 | 48 | 13 | DG | PM-p2 |
| 2 | BCC6 | 202550 | NLI-IF phosphatase* | -0.39 | 233 | 2212 | 39 | 4 | IMC | IMC |
| 1 | BCC7 | 311230 | hypothetical; 1 TM domain | 0.74 | 495 | 4603 | 202 | 11 | IMC | IMC |
| 1 | BCC8 | 273050 | hypothetical | 2.08 | 71 | 642 | 0 | 2 | Golgi | PM-p2 |
| 4 | BCC9 | 200330 | hypothetical | 0.26 | 67 | 589 | 0 | 12 | N/A | N/A |
| 3 | BCC10 | 310220 | guanylate-binding/atlastin | -1.36 | 150 | 1396 | 46 | 10 | N/A | N/A |
| 3 | BCC11 | 278130 | hypothetical | -0.10 | 168 | 1534 | 22 | 14 | chromatin | PM-p2 |

### Assembly of the BC map

To further expand the list of BC components and refine its sub-structural architecture we expanded the BioID bait set with new BC residents BCC1 and BCC2, as well as previously characterized myosin J (MyoJ), which temporal localization to the BC is similar to Cen2 (MyoJ; (Frenal et al., 2017)). The collective mass spectrometry data were used to generate prey-prey maps by correlating the abundance of individual preys over all biotinylation experiments, and thus mapping potential colocalization and thereby visualizing subcellular complexes (Knight et al., 2017; Youn et al., 2018) (Fig 2a, Fig. S2a). In addition, the data set was probed for more detailed statistical support for reciprocal discoveries by using the Significance Analysis of INTeractome (SAINT) algorithm (Choi et al., 2011; Teo et al., 2014), designed to identify interaction partners in AP/MS or proximity-biotinylation applications (Fig 2b, Fig. S2b, Table S1). To assure we were capturing as much as possible of the BC proteome we analyzed the data with a variety of different settings. We also ran the analyses with or without the inclusion of a cytoplasmic BioID2-YFP fusion control (Engelberg et al., 2020) to reduce the false positive detection of spurious cytoplasmic proteins randomly interacting with the BC-localizing BioID2 baits. The goal of this wide-cast trial-and-error approach was to refine the settings define the cut-off for a guilty-by-association strategy to predict which candidates are novel BC proteins. Throughout this process we tagged putative BCC proteins, which besides BC proteins mapped the apical annuli (Engelberg et al., 2020), as well as several new proteins co-localizing with Cen2 at the apical end and proteins in the nuclear centrocone with MORN1 (Table S1). Not a single setting provided a clear-cut definition suggesting significant overlap between the BC and other compartments, which in part stems from the multiple sub-cellular structures that both MORN1 and Cen2 are associated with, as well as the feature that proteins deposited in the daughter cytoskeleton during division are recruited at the BC but upon maturation end up in other non-BC cytoskeleton structures like the apical annuli and the alveolar sutures. The tightest analysis representing comprising all newly mapped BCCs is provided in the prey-prey map of Fig. 2a, whereas select details of the statistical analysis resolved per bait are provided in Fig. 2b, whereas more extensive analyses are included in Fig. S2 and Table S1.

**Fig 2.**
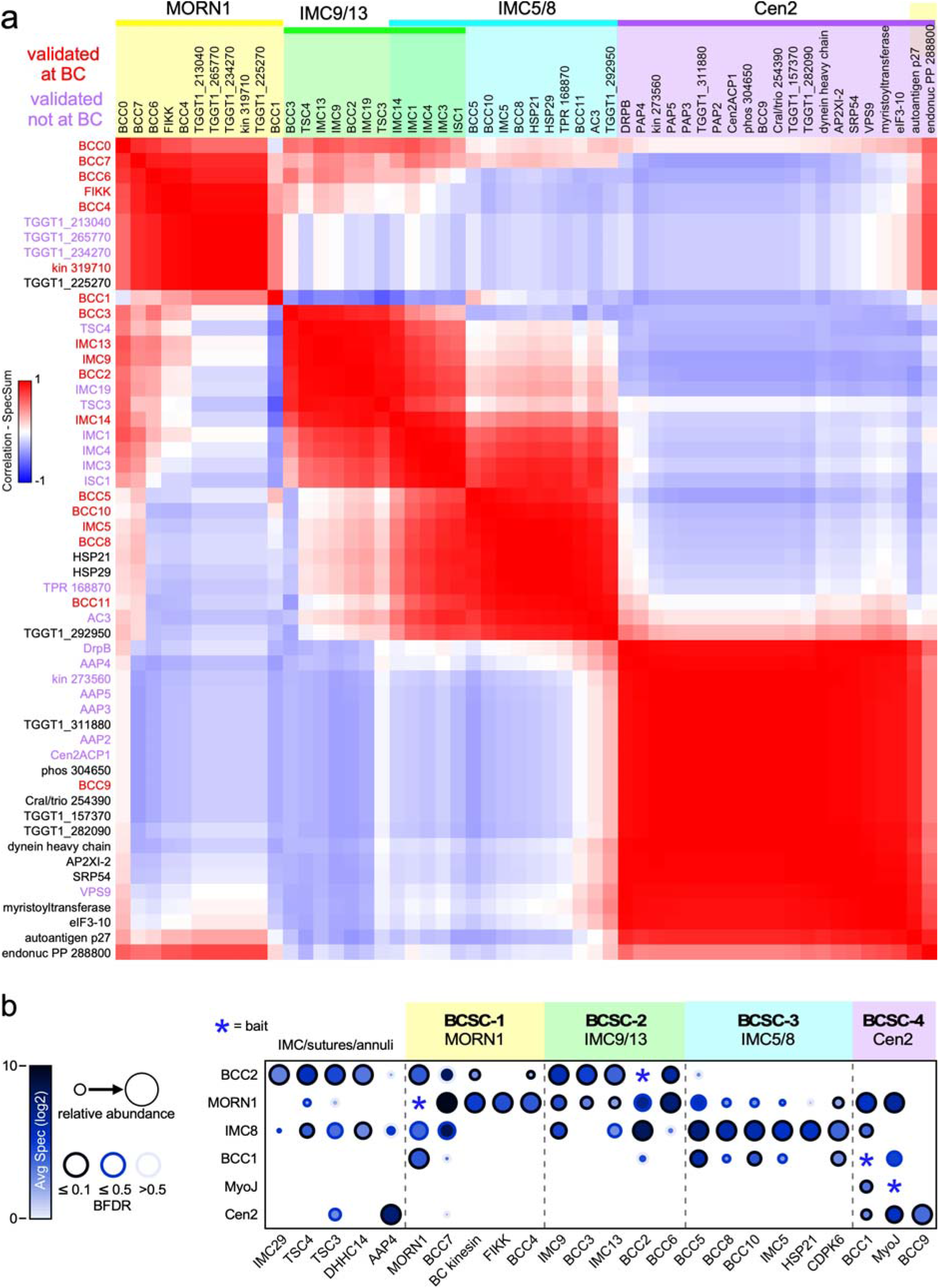
Reconstruction of the BC architecture via spatial relation analysis. **a.** Distance heatmap visualizing the BC proximity landscape, which partitioning into four major BC sub-complexes (BCSCs) marked in yellow (BCSC-1), green (BCSC-1) (BCSC-1), blue (BCSC-1) and purple (BCSC-1). Proteins with overlapping partitioning to two different BCSCs are dual-colored accordingly. The defining and localization validated components are named at the top of each BCSC defining, colored group. The average of the spectral counts for a given prey were correlated to every prey that passed the threshold of FDR 0.1 and visualized in the distance heatmap. Prey order is the same for the x- and the y-axis. See Suppl Fig S1 for inclusion of a different control resulting in loser cluster definition. **b.** Dotplot summarizing known and newly identified BCCs over the executed experiments, including the BC proteins used as BioID2 fusion baits. Preys that pass a threshold of FDR < 0.5 were visualized and manually sorted in the four BCSCs as defined in panel A (BCSC-1 assignments were based on panel ‘a’ as well as additional analyses using different settings and controls; see Suppl Fig S1. The IMC/sutures/annuli group defines a separate group in this analysis, which shares overlap connections mostly with BCSC-2 and to a lesser extend with BCSC-3. The entire dotplot is shown in Fig. S2B.

The prey-prey maps and SAINT analyses provided complementary, deep insights into the BC architecture. The prey-prey distance heat maps consistently revealed four major clusters, which we color coded and named BC Sub-Clusters 1-4 (BCSC1-4; Fig. 2a, Fig. S2a). These clusters confirmed our previous observations of the BC architecture that were anchored on MORN1(BCSC-1), IMC9/13 (BCSC-2), IMC5/8 (BCSC-3) and Cen2 (BCSC-4) (Anderson-White et al., 2011). However, as noted above, several additional subcellular localizations besides the BC were interspersed in these clusters, including the serendipitous discover of a set of apical annuli proteins (AAPs) for Cen2 (Engelberg et al., 2020) and centrocone proteins (e.g. TGGT1_213040, TGGT1_265770, TGGT1_258450, and TGGT1_270810) for MORN1. Therefore, assign uncharacterized components in these clusters to the BC substructures through ‘guilty-by-association’ strategy in the prey-prey maps was not reliable, but became a lot more refined by adding the SAINT analyses represented in the dot-plots (Fig. 2b, S2b). Especially candidates exclusively detected with Cen2 or MORN1 could in general be triaged for presence in the BC (Fig. S2b). With this in mind, we can with reasonable confidence assign hypothetical protein TGGT1_225270 to BCSC-1, and CDPK6 and HSP21 to BCSC-3 without experimental validation. Altogether, we validated a total of 11 new BCC proteins, distributed over the 4 BCSCs (Table 1, Fig. S1).

In addition, we find a cluster of proteins not present in the BC, but which provides insights in how the BC is interfacing with the IMC, e.g. IMC29 and suture proteins TSC3 and 4, (Chen et al., 2017), or is the site of growth of the daughter (e.g. palmitoyl transferase DHHC14, which is critical for IMC assembly (Dogga and Frenal, 2020)) (Fig. 2b, Table S1). Whether these hits represent direct connection between the BC and the IMC or whether these are hits simply due to their BC proximity can not be differentiated using the current data, but the detection of this set of protein across 2-4 BC baits, notably IMC8, suggests these interactions are real (Fig. 2b, S2b). A scenario where for example AAP4 is first recruited to the BC before being deposited in the annuli is conceivable, and as such might provide insights in how the BC is involved in daughter bud assembly. Overall, reciprocal BioID of BC components aligned with the three BC substructures previously mapped, identifies surprising associations between the BC and IMC structures, and assigns several previously uncharacterized hypothetical proteins to different BC sub-structures.

### The novel BCCs comprise a diverse set of proteins

We characterized the new BCCs for putative function by mining ToxoDB (Gajria et al., 2008) for the following data: lytic cycle fitness score (Sidik et al., 2016), phosphoproteome (Treeck et al., 2011), hyperLOPIT subcellular localization assignments (Barylyuk et al., 2020), size, number of exons, and functional (domain) annotation (Table 1). This revealed that 8 BCCs are hypothetical proteins, with no known domains or function. However, BCC2 contains a kinase domain, BCC5 and BCC6 are phosphatases, but none of these have fitness scores under -2, which is in general a good indicator of potential essential roles in the lytic cycle (Sidik et al., 2016) and might indicate redundant functions. Phosphatase BCC5 further harbors an EF-hand, indicating this protein likely binds calcium, whereas BCC6 is annotated as a nuclear LIM interacting factor (NLI-IF) family phosphatase. NLI phosphates typically dephosphorylate the RNA Polymerase II C-terminal domain, which regulates their activation. However, such domain is also found in many *Arabidopsis thaliana* phosphatases, which have much more divergent functions (Bang et al., 2006). BCC10 harbors a guanylate-binding domain and a weak homology to atlastin, which in vertebrates is a dynamin-like GTPase required for homotypic fusion of endoplasmic membrane tubules (Betancourt-Solis et al., 2018). However, the mild fitness score indicated BCC10 is not essential. Finally, BCC7 harbors a TM domain, whereas BCC0 contains predicted myristoylationa and palmitoylation sites at its N-terminus, which makes these the only two BCCs with a likely membrane anchoring. Of all BCCs, only BCC0 and BCC4, and maybe BCC1, had fitness scores indicating a potential essential role, which prioritized them for further functional characterization.

Furthermore, we performed a global analysis of distinct BC spatio-temporal localization kinetics for the various BCCs (Fig. S1). This revealed that BCC6 and 7 exclusively localize to the BC of mature parasites, following completion of cell division (like FIKK (Skariah et al., 2016) and MSC1a (Lorestani et al., 2012)), whereas BCC 3 and 4 exclusively localize to the budding daughter BC, however, BCC3 presents a special case as it localizes to the whole daughter bud, which prioritized it for follow up. All other BCCs (1, 2, 5, 8, 9, 10, 11) localize to the BC of both the budding daughters and mature mother. Combining all criteria, we selected BCC0, 1, 3, and 4 for more detailed experimental exploration, which we will present below in order of appearance at the BC.

### BCC0 provides a foundation for the BC and other cytoskeletal elements

The severe -4.10 fitness score suggests BCC0 is essential for completing the lytic cycle. Besides a predicted myristylation site and three palmitoylation sites the primary structure does not provide clues toward its function (Fig. 3a). C-terminal tagging of the BCC0 locus with a spaghetti monster (sm)-Myc-tag revealed its association with the centrosomes very shortly after their duplication and along the process of centrosome separation condenses in several foci surrounding the centrosomes (Fig. 3b). To assess whether BCC0 is recruited before ISP1, a palmitoylated IMC component that is one of the earliest scaffold markers (Beck et al., 2010), we co-stained with ISP1 antiserum. Indeed, BCC0 is detected well before ISP1 is recruited to the daughter bud, making it one of the earlier known markers of the daughter cytoskeleton (Fig. 3c). ISP1 first shows up with a moderate intensity before it extends across the apical cap alveolus and becomes more intense (Beck et al., 2010), at which point BCC0 has already started to extend in the basal direction away from ISP1 in a stringy pattern of spots reminiscent of the longitudinal sutures between the alveolar plates (Chen et al., 2015a; Lentini et al., 2015; Chen et al., 2017). Since none of these patterns are consistent with a BC localization, we visualized MORN1 with an endogenous YFP fusion and performed SR-SIM super-resolution microscopy. This strikingly revealed a very defined pattern of in general 5 BCC0 foci laying right on top of the early BC (Fig. 3d). The early timing in daughter development is communicated by the barely separated MORN1 signals in the centrocones, which mark the spindle poles. Note that the 5 foci are a consistent observation and can also be appreciated in Fig. 3c. Next we asked whether these 5 foci could represent (the basis) of the apical annuli, which display a similar five-fold symmetry at the suture separating the apical cap from the rest of the IMC (Engelberg et al., 2020). We interrogated this by co-staining BCC0 with AAP4 antiserum. Neither during initiation nor at a later time point of development could we appreciate significant co-localization of AAP4 and BCC0 (Fig. 3d, Supplementary Movie 1; note that the AAP4 anti-serum aspecifically cross-reacts with the centrosome (Engelberg et al., 2020)). This suggests they might not be structurally related, although we cannot exclude the possibility that BCC0 provides the foundation for the annuli, which do not appear till later in cell division (Engelberg et al., 2020) at which point BCC0 might already have extended along the longitudinal sutures. Overall, BCC0 seems to lay down an early 5-fold symmetry transitioning into the sutures, and positionally is set to form a potential basis for establishing the BC. It is this BC preceding appearance that led to naming this protein BCC0, deviating from our naming convention mentioned earlier.

**Fig 3.**
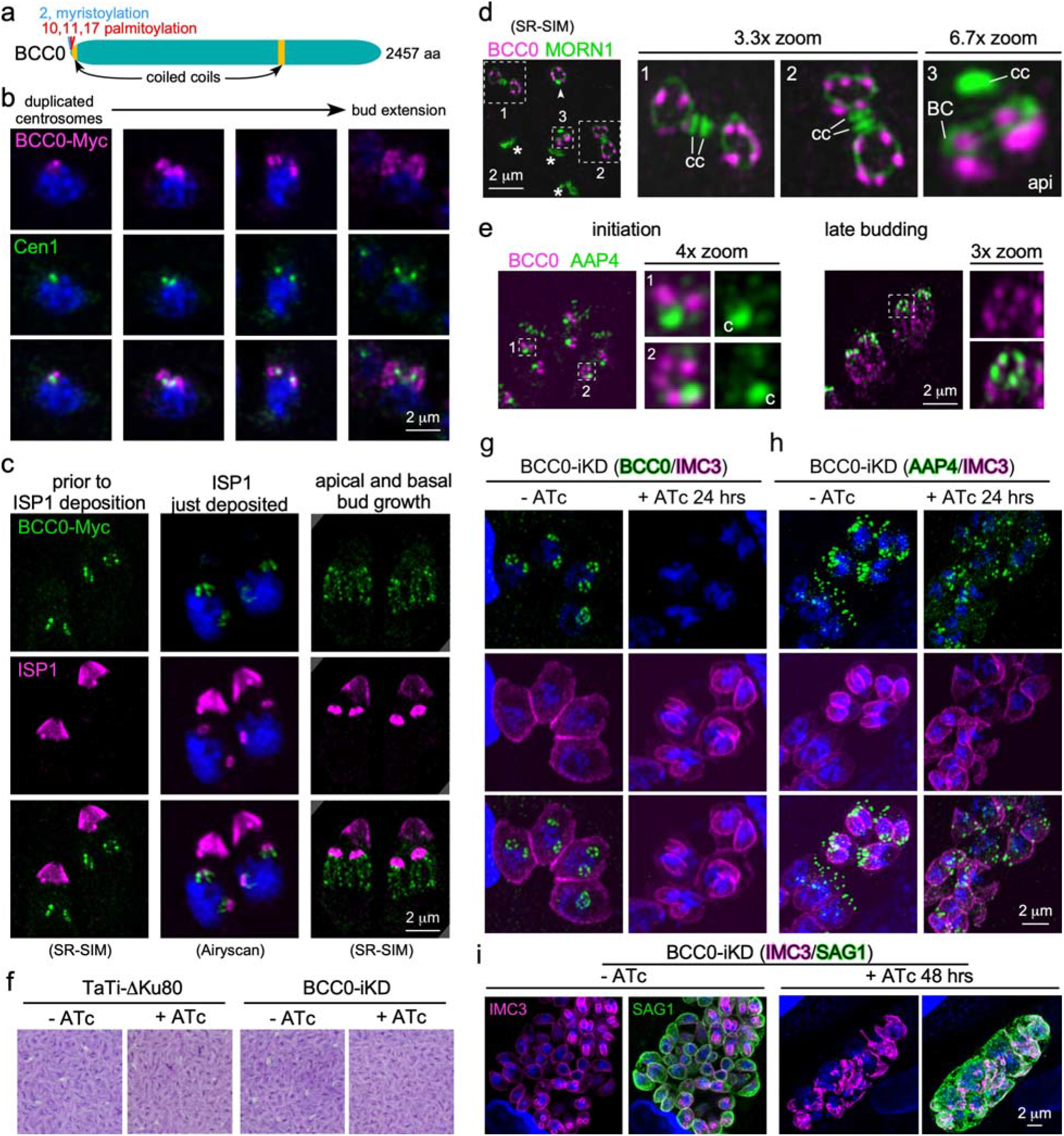
BCC0 is a novel component of the nascent daughter bud scaffold. **a.** Schematic of the protein. BCC0 harbors a G in position two suggesting putative myristylation and contains three C residues for putative acylation at its N-terminus, next to two coiled-coil domains, as the only appreciable annotations. **b.** Parasites wherein BCC0 was endogenously tagged with the spaghetti monster Myc tag (smMyc) were co-stained with antiserum against human Centrin2 (marking the *T. gondii* centrosome outer core) and demonstrate that BCC0 assembles early in division around the just duplicated centrosomes and as daughter buds grow, BCC0 highlights the nascent bud scaffold in a spotty pattern. DNA is stained with DAPI (blue). **c.** BCC0-smMyc colocalization with ISP1 antiserum reveals that BCC0 is recruited to the daughter bud earlier in the division cycle than ISP1 and localizes basal to the ISP1 signal. When ISP1 is deposited, BCC0 starts to elongate in the basal direction and has the appearance of the longitudinal IMC sutures in larger daughter buds. **d.** SR-SIM of BCC0-smMyc co-expressing endogenously tagged YFP-MORN1 shows that the BCC0 signal at the time of MORN1 appearance at the BC is assembled as 5-6 dots apical to the BC. Boxed regions numbered 1-3 in the left panel are magnified in the corresponding panels on the right. With daughter bud maturation, BCC0 highlights the longitudinal sutures of the nascent daughter scaffold. Asterisks mark the BC of the mother; arrowhead marks the other centrosome and daughter bud in the middle parasite corresponding with box #3; ‘c’ marks the centrocones (spindle poles); ‘api’ marks the apical end of this particular bud. DNA is stained with DAPI (blue). A 3D rendered rotation of this parasite is provided in Supplementary Movie S1. **e.** Airyscan images of BCC0-smMyc parasites co-stained with AAP4 antiserum. Note that at neither stage of division, the BCC0 foci co-localize with apical annuli serum but appear to form slightly distinct constellation clusters (it remains possible that AAP4 is recruited to the 5 BCC0 foci, at which point they disperse into the sutures). **f.** Plaque assays of inducible knock-down of BCC0 (BCC0-iKD). The endogenous promoter of BCC0 was replaced with a TetO7sag4 anhydrous tetracycline (ATc) regulatable promoter, demonstrating that BCC0 is essential for the lytic cycle. TaTi-ΔKu80 is the parent line. Plaque assays performed for 7 days. **g.** BCC0-iKD parasites co-stained with Ty and IMC3 antisera, highlighting BCC0 and the IMC cytoskeleton scaffolds, respectively. BCC0 depletion results in disorganized daughter IMC and nuclear partitioning defects. DNA is stained with DAPI (blue) **h.** BCC0-iKD parasites co-stained with AAP4 and IMC3 antisera demonstrates that BCC0 depletion does not disrupt the AAP4 annuli foci, but they become misorganized consistent with the IMC biogenesis defects. DNA is stained with DAPI (blue) **i.** BCC0-iKD parasites co-stained with SAG1 and IMC3 antisera highlights that the parasites continue to expand their size even after 48 hrs ATc treatment, but they lose their typical banana shape, marked by plasma membrane (PM) marker SAG1. The misorganized IMC no longer encases the parasite nuclei and appears only sparsely associated with the PM.

To dissect BCC0 function we replaced its promoter with a tetracycline regulatable promoter (Meissner et al., 2002). Plaque assays confirmed that BCC0 is essential for completing the lytic cycle (Fig. 3f) and BCC0 indeed depleted upon ATc addition (Fig. 3g). BCC0 depletion leads to aberrant IMC structures, which have a very wide appearance, and some nuclei are not encased by IMC (Fig. 3g-i). AAP4 still assembles in foci on the IMC suggesting that BCC0 might not be critical for apical annuli assembly. Furthermore, prolonged BCC0 depletion results in a large cytoplasmic mass (visualized with plasma membrane marker SAG1; Fig. 3i). Overall, even though BCC0 is never physically present at the BC, the phenotype indicated that BCC0 is important for correct IMC3 assembly and association with the plasma membrane where the wide appearance of the IMC is highly suggestive of BC involvement.

### BCC3 dynamically localizes to the bud initiation complex, the sutures, and the BC

BCC3 was selected for its absence from the mother BC and its exclusive association with the daughter cytoskeleton. Besides a coiled coil domain no functional features can be appreciated in BCC3 (Fig. 4a). We tagged BCC3 at the C-terminus with a triple Myc-tag and imaged it in detail throughout parasite division using IMC3 as bud marker as it had a very dynamic appearance. We immediately noticed that BCC3 also appears as 5 foci early in cell division just before IMC3 shows up at the daughter scaffold. Throughout daughter budding, BCC3 sequentially is seen in a pattern reminiscent of the longitudinal sutures, whereas mid budding an addition signal condensation appears to form at the suture below the apical cap, while toward the end of budding the signal exclusively localizes to the BC of the daughters before completely disappearing upon conclusion of cell division (Fig. 4b). Co-staining with YFP-tagged MORN1 showed that besides co-localizing the BC, at the other end of the bud BCC3 sits right below MORN1 at the apical end of the IMC (Fig. 4c, blue arrowheads). Co-staining with AAP4 showed that BCC3 localization was bordered by the apical annuli and did not extend into the apical cap. Early in division, the apical cap still has a very flat appearance, suggesting it is extending later than the IMC assembling in the basal direction (Beck et al., 2010). Since BCC3 has a modest fitness score (- 0.11) we did not attempt a gene knock-out as it is unlikely to have a critical role. In conclusion, BCC3 is a very dynamic daughter bud marker present at its foundation as 5 foci and then transitions onto the daughter scaffold sutures before accumulating on the BC in the conclusion of cell division and is completely released from the cytoskeleton upon maturation.

**Fig 4.**
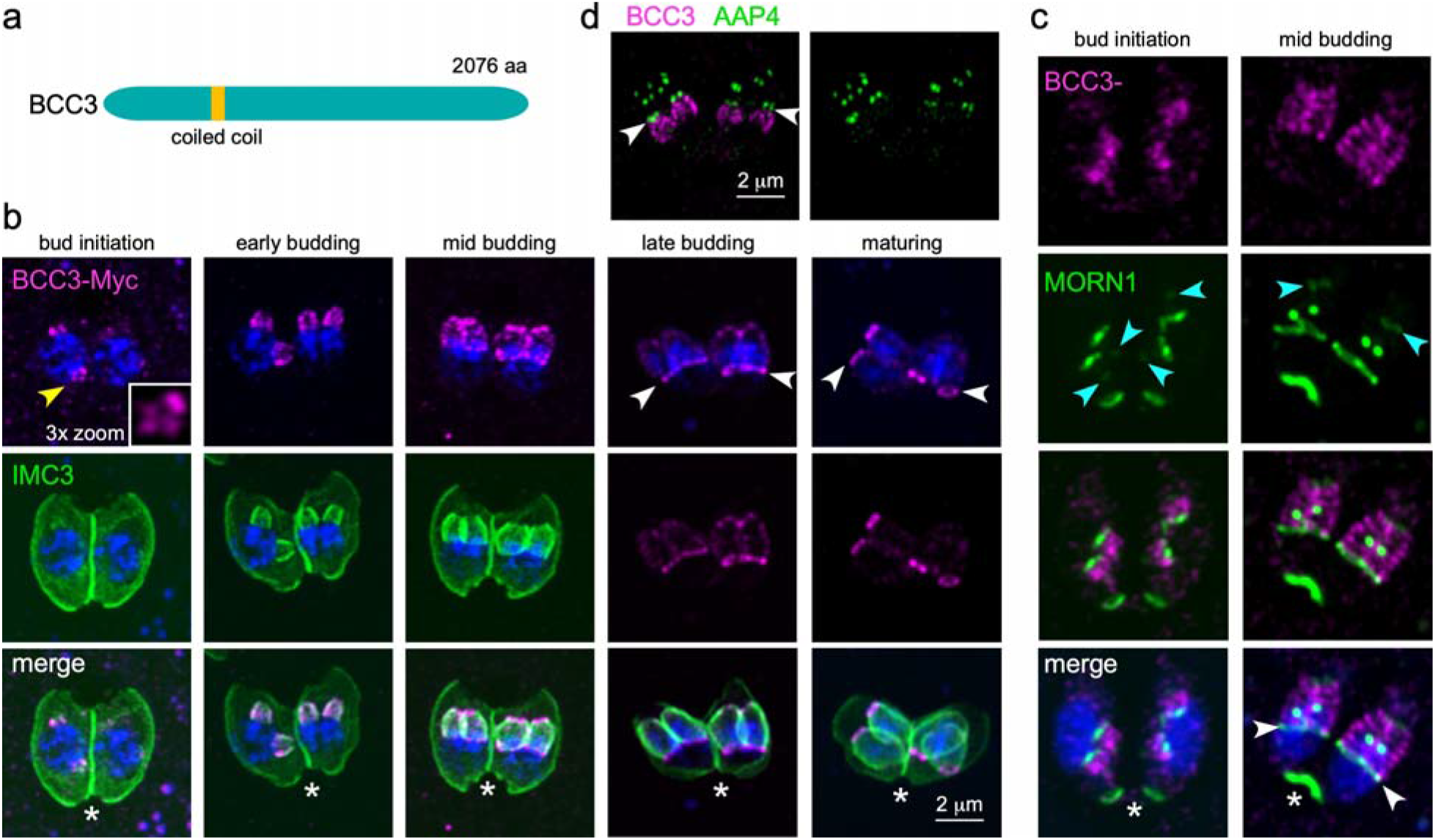
BCC3 dynamically localizes on the IMC scaffolds and BC during daughter budding. **a.** Schematic representation of BCC4, which only harbors a single coiled-coil region as recognizable domain. **b.** Parasites expressing endogenously C-terminally Myc3-tagged BCC3 were co-stained with IMC3 antiserum and shows that BCC3 localizes to the 5-foci early in cell division (yellow arrowhead, enlarged in box) and then extends basally along the developing cytoskeleton in a speckled pattern through the first half of budding. Upon maturation of the daughters, BCC3 transitions completely to the daughter BC (arrowheads) and eventually releases from the cytoskeleton in mature parasites (asterisks). DNA is stained with DAPI (blue) **c.** Co-stain of Myc3-tagged BCC3 with endogenously tagged YFP-MORN1. Blue arrowheads mark MORN1 at the apical end of the IMC, and the BCC3 signal extends almost to this signal in the early stages, but at mid budding BCC3 ends more basally in a thick line, likely at the suture below the apical cap (see the same stage in panel b for a comparable signal). Note the localization of BCC3 to the daughter bud IMC sutures during the mid-steps of division and the prominent co-localization with MORN1 in the daughter BC later in division (white arrowheads) but BCC3 is absent from the BC of the mother cell (asterisks). Scale bars are the same as shown in panel b. DNA is stained with DAPI (blue). **d.** Co-stain of 3xMyc-tagged BCC3 with AAP4 antiserum to mark the apical annuli in an early-mid stage budding parasite demonstrates that BCC3 is not present in the apical cap and suggests the apical cap grows apically at the same time as the more basal alveoli grow in the basal direction.

### BCC4 exclusively associates with the budding daughter BC

BCC4 stood out for its severe fitness score (−3.52) in the lytic cycle screen, suggesting it is an essential protein (Sidik et al., 2016). BCC4 is a hypothetical protein harboring extensive low complexity regions and comprises a single coiled-coil domain, but has otherwise no identifiable domains (Fig. 5a). We tracked endogenously BCC4 tagged with 3xMyc in detail throughout cell division using a β-tubulin co-stain as guide. BCC4 appears early in the division cycle in close proximity to the spindle microtubules and are associated with early daughter bud formation (Fig. 5b). For the remainder of division BCC4 is present at the BC of nascent buds, but upon emergence of daughter cells from the mother the BCC4 signal disappears. Since BCC4 does not localize to the BC in mature parasites, this supports an exclusive and essential function during cell division. To further pinpoint the early temporal events, we colocalized BCC4 with a marker of the centrosome outer core from which bud formation nucleates. BCC4 accumulated on top of the just duplicated centrosomes (Fig 5c, 1), and forms rings around the centrosomes while S/M-phase progresses (Fig 5c, 2-3). Since the early BC formation is mediated by MORN1, which also forms rings around the dividing centrosome (Gubbels et al., 2006; Hu et al., 2006; Hu, 2008; Suvorova et al., 2015), we co-colocalized MORN1, BCC4 and the centrosome. This demonstrated that BCC4 and MORN1 display the exact same dynamics during BC formation: both proteins initially accumulate distal to the centrosome (Fig 5d) and subsequently transform into the ring-shaped BC following centrosome duplication (Fig 5e). Thus, BCC4 and MORN1 are recruited to the nascent BC with similar kinetics. However, in contrast to MORN1, BCC4 exclusively localizes to the BC and is released from the BC when cell division completes (Fig 5e, arrowheads), thereby sharply focusing its role in cell division.

**Fig 5.**
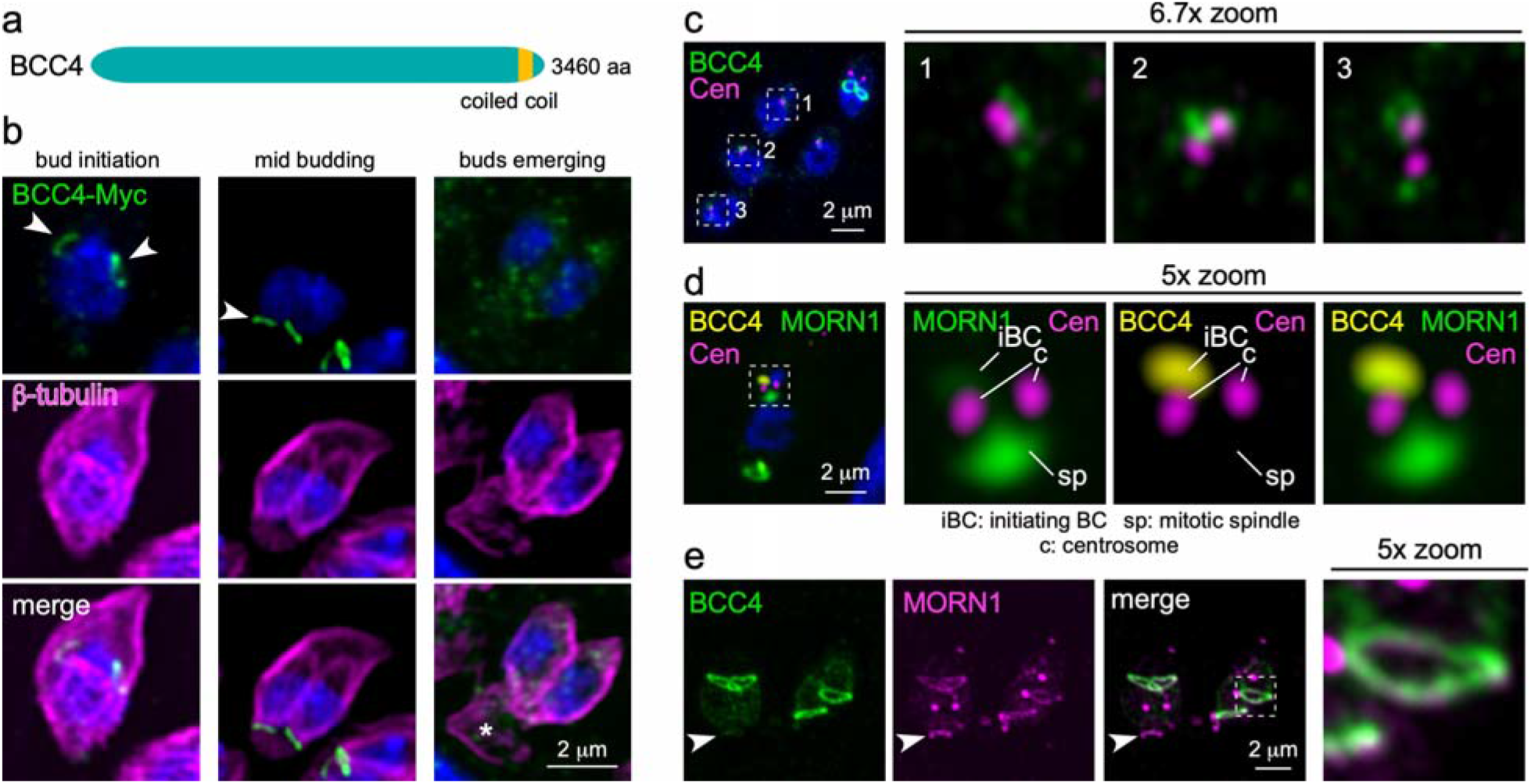
BCC4 only localizes to the BC during division. **a.** Schematic representation of BCC4, which only harbors a single coiled-coil region as recognizable domains. **b.** Parasites expressing endogenously C-terminally Myc3-tagged BCC4 were co-stained with -β- tubulin antiserum and shows that BCC4 localizes to the BC right at its formation but is absent from the mature BC. Arrowheads mark the early-and mid-development daughter BCs; asterisk marks the retracting and disassembling mother’s cytoskeleton. DNA is stained with DAPI (blue). **c.** Co-staining of BCC4-Myc parasites and the *T. gondii* centrosome (*Hs*Centrin2 antiserum) shows that BCC4 assembles as a ring-like structure at the point of centrosome duplication. The boxed and numbered panels are magnified as indicated and show BCC4 transitioning from an undefined bleb around the centrosome in panels 1 and 2 into a ring as visible in panel 3. DNA is stained with DAPI (blue). **d.** Four color imaging displays the interplay of YFP-MORN1 and BCC4-Myc3 and the centrosomes in early BC formation. At this very early stage of daughter development before completion of spindle pole separation, BCC4 and MORN1 co-localize in an amorphous mass in close apposition to the outer centrosome. DNA is stained with DAPI (blue). *T. gondii* **e.** YFP-BCC4 colocalizes with 5xV5-MORN1 in the BC around the midpoint of daughter cell budding, but in contrast to MORN1, BCC4 is absent from the mature BC in the mother parasite (arrowhead).

### BCC4 is an essential BC component

To test the function of BCC4 we replaced its promoter with a tetracycline-regulatable promoter. Plaque assays for 7 and 14 days display no plaques in the presence of ATc and demonstrate that BCC4 is essential for *in vitro* proliferation (Fig. 6a). Phenotype analysis by IFA using AAP4 and IMC3 markers showed that depletion of BCC4 results in the formation of double headed parasites conjoined at their basal end (Fig. 6b). The same phenotype was also seen upon depletion of MORN1, which was the result of a defect in BC assembly (Lorestani et al., 2010), as well as upon depletion of phosphatase HAD2a (Engelberg et al., 2016).

**Fig 6.**
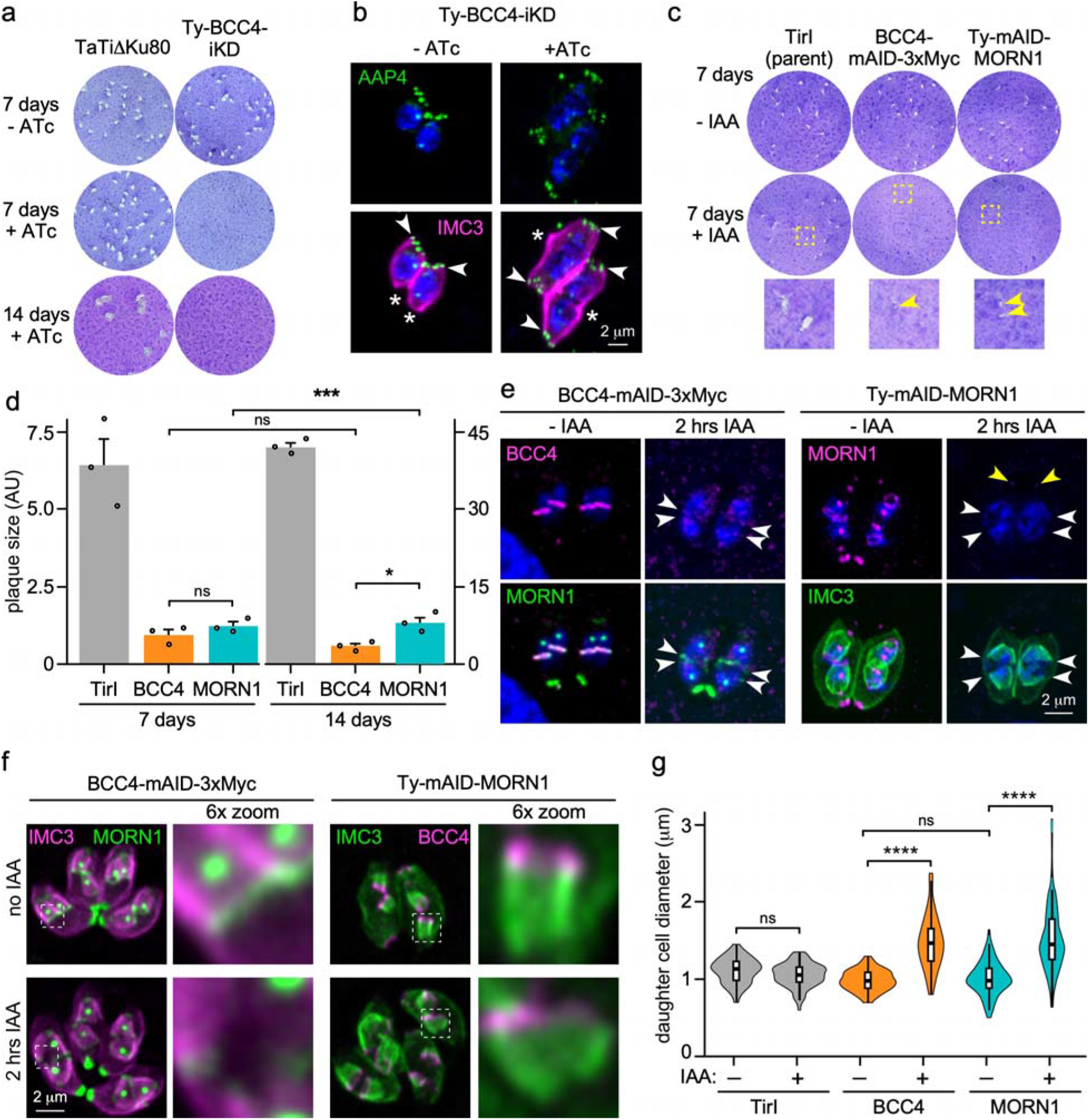
BCC4 is essential for completing the lytic cycle by way of BC constriction. **a.** Replacement of the BCC4 promoter with a TetO7sag4 anhydrous tetracycline (ATc) regulatable promoter completely abolishing plaque forming capacity and demonstrates BCC4 is essential to complete the lytic cycle. At the same time as promoter replacement, an N-terminal Ty-tag was introduced on the BCC4 ORF. **b.** BCC4 knock-down for 24 hrs with ATc results in the formation of double headed daughters. AAP4 antiserum stains the apical annuli and indicates the apical end (arrowheads), whereas the basal end of the parasite is marked with asterisks. IMC3 antiserum marks the cortical cytoskeleton. DNA is stained with DAPI (blue). **c.** Plaque assays of parasites wherein either BCC4 or MORN1 is fused with the mini auxin inducible degron (mAID) sequence to permit rapid protein degradation upon addition of auxin (indole-3-acetic acid or IAA) demonstrate the presence of small plaques in both cell lines after 7 days of IAA treatment. This illustrates limited or severely reduced parasite proliferation. **d.** Quantification of plaque size after 7 and 14 days of IAA treatment highlights that BCC4 stops proliferating (no significant plaque size increase between day 7 and 14), whereas MORN1 degradation results in a continuous but severely reduced parasite proliferation rate (significant increase in plaque size between day 7 and 14). 7 day scale on the left-axis; 14 day scale on the right-axis. TirI parasites are the parent line and serve as wild type control. Cell lines grown without IAA were included and counted, but not included in the depicted plot. n=3 + sem. Statistical significance was tested with one-way analysis of variance (ANOVA; F_11_,_24_=461.8, p=2^-16^) followed by post hoc Tukey’s test to determine significance. **e.** Fast kinetics of mAID mediated knock-down of BCC4 and MORN1 for 2 hrs. BCC4 is completely gone from the BC visualized by endogenously expressed YFP-MORN1 that still marks the BC (arrowheads). 2 hr induction of Ty-mAID-MORN1 parasites (co-stained with IMC3 antiserum) results in depletion of MORN1 from the centrocone and BC (white arrowheads), but not from the apical end (yellow arrowheads). DNA is stained with DAPI (blue). **f.** BCC4 and MORN1 mAID parasites reciprocally endogenously expressing YFP=MORN1 or 3xMyc-BCC4 to report on the BC were treated with IAA for 2 hrs and co-stained with IMC3 antiserum. Both MORN1 and BCC4 knock-down parasite display wider basal gaps of the daughter buds compared to untreated parasites. Dotted line boxes marking representative basal daughter ends in the left side panels are magnified in the right panels as indicated. **g.** Violin plots of quantified daughter basal complex opening defects shown in panel ‘f’. TirI parasites are the parent line and serve as wild type control. Each condition represents at least 50 daughter basal ends measured; the whisker bar indicates the median, the whisker box upper corner resembles 75^th^ percentile and the lower whisker box corner the 25^th^ percentile. Whiskers mark highest and lowest value measured, respectively. Significance was tested with one-way ANOVA (F_5,426_=60.44, *p*=2^-16^) followed by post hoc Tukey’s test to determine significance. ns= not significant, **** *p*<0.0001.

Since promoter-replacement acts on gene transcription and relatively slowly permeate into protein kinetics, we also fused BCC4 to the auxin-inducible degron (AID) system to deplete BCC4 at a much faster rate (Brown et al., 2017). We further generated a cell line in which MORN1 was tagged at its 5’end with Ty-mAID to directly compare phenotypic consequence between the two BC components. Plaque assays with either BCC4-mAID-3xMyc or Ty-mAID-MORN1 parasites exhibited small plaques after 7 days of IAA treatment (Fig. 6c). Notably, plaques were found to be ∼7-times smaller compared to the parental or non-induced populations, which indicates parasite proliferation is severely inhibited although not completely disrupted in the AID system (Fig. 6d). Interestingly, after prolonged 14 days of growth the size of Ty-mAID-MORN1 induced plaques significantly increases, which is not seen for the BCC4-mAID auxin-treated parasites (Fig. 6d). For both the MORN1 and BCC4 mAID lines, the proteins were largely depleted after two hours of IAA treatment, which already resulted in wider basal ends of the parasite buds (Fig. 6e). To follow BC fate after BCC4 or MORN1 degradation, we also tagged BCC4 and MORN1 reciprocally in the corresponding MORN1 and BCC4 mAID parasite lines. This showed that 2 hrs of MORN1 depletion does not have an impact on BCC4 recruitment to the daughter BC, and vice versa (Fig. 6f). However, we again observed a wider daughter basal end in either scenario with daughter cells appearing almost bell-shaped instead of their narrow appearance seen in the control or not induced lines. Measurements of the daughter basal end diameters showed the widths were similar in the BCC4 and MORN1 depleted parasites, which were significantly larger than detected in wild type or non-induced controls (Fig. 6g). Collectively, these data indicate that BCC4 depletion phenocopies MORN1 depletion.

### BCC4 is required for maintaining BC integrity throughout cell division

We next analyzed phenotypic consequences of 24 hrs BCC4 depletion, which induced the same double-headed phenotype as seen with the ATc conditional system (Fig. 7a vs Fig. 6b). To ascertain these double-headed parasites indeed have two apical ends we co-stained with AAP4 antiserum. Both BCC4-mAID_3xMyc and Ty-mAID-MORN1 parasites treated with IAA for 24 hrs exhibited AAP4 staining at both ends of the double-headed parasites (Fig. 7b). Quantification of the number of double-headed parasites per vacuole revealed a significant difference between MORN1 and BCC4: BCC4 depleted parasites show a nearly 2-fold higher incidence of more than two double-headed daughter parasites per vacuole (Fig. 7c). This indicates that BCC4 depletion more profoundly affects parasite division, leading to a more penetrant phenotype consistent with the results from the plaque assays (Fig. 6c).

**Fig 7.**
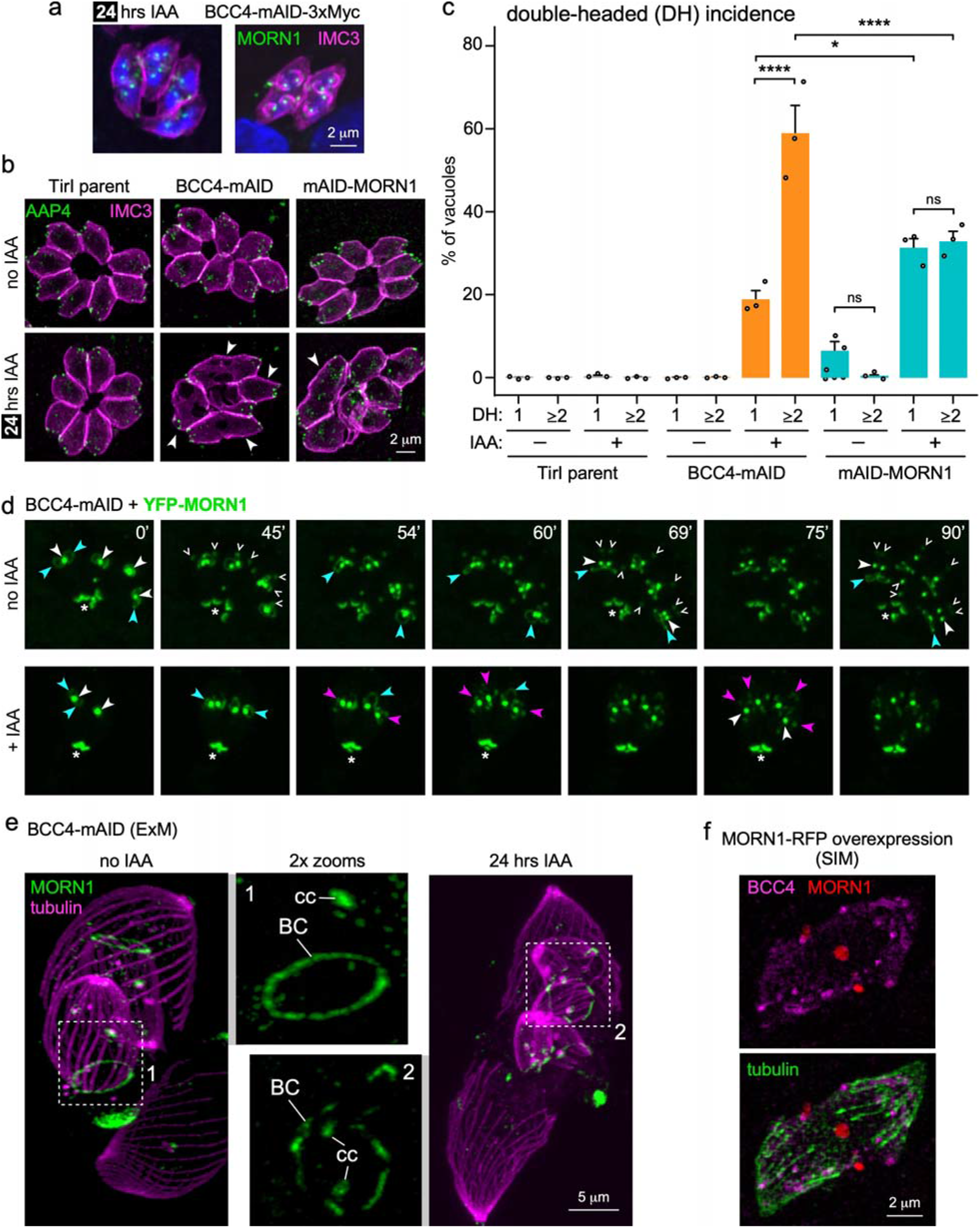
BCC4 and MORN1 are both needed to maintain BC integrity during endodyogeny. **a.** BCC4-mAID parasites endogenously expressing YFP-MORN1 were treated with IAA for 24 hrs and co-stained with IMC3 antiserum. The left panel shows parasites that did not complete cell division and harbor two nuclei with two MORN1 centrocone foci per nucleus indicating they started a next round of cell division. The right panel shows incompletely divided parasites that are undergoing the next round of daughter budding, with four daughter buds per parasites per mother cell, each associated with a MORN1 centrocone (spindle pole) in the nucleus indicating there are no defects in the nuclear cycle. Note the bell-shape appearance of the daughter buds due to loss of basal integrity. DNA is stained with DAPI (blue). **b.** BCC4-mAID and Ty-mAID-MORN1 parasites treated with IAA for 24 hrs are capable to initiate rounds of intracellular division, but fail to conclude division and exhibit a “double-headed” phenotype (arrowheads), displaying two apical ends (highlighted by AAP4 staining) located at the extreme ends of conjoined parasites. Note the difference of “double-headed” frequency per vacuole between BCC4 and MORN1 knock-down parasites. **c.** Quantification of “double-headed” phenotypes in BCC4 and MORN1 IAA treated parasites. Parasites were incubated for 24 hrs in IAA before fixation and staining with IMC3 and AAP4 antisera. 100 random vacuoles were quantified for inhabitation by one “double-headed” parasite (1 DH) and vacuoles that inhabit more than one “double-headed” parasite (≥ 2DH). Average of n=3 biological replicates, +sem is shown. Significance tested with one-way ANOVA (F_11,24_=69.68, p=1.85^-15^) with post hoc Tukey test. ns= not significant, * *p* <0.05, ** *p* <0.01, *** *p* <0.001, **** *p* <0.0001. **d.** Select time-lapse microscopy panels of BCC4mAID-3xMyc parasites expressing endogenously YFP-tagged MORN1. IAA was added 2 hrs before t = 0 min. Although BCs initially form normally upon BCC4 depletion, they start falling apart (scattered MORN1 at the BC) at t = 54 min, and all BCs are fragmented by t = 69 min. MORN1 localization in select panels is annotated as follows: open arrowheads, apical end of budding daughters; closed white arrowhead, centrocone (spindle pole); bleu arrowhead, BC of budding daughters; magenta arrowhead, fragmented BC of budding daughters. Panels correspond with Supplementary Movies S2 and S3. **e.** Expansion microscopy (ExM) of BCC4-mAID-3xMyc parasites endogenously expressing YFP-MORN1 co-stained with α-tubulin (12G10) antiserum. BCC4 depletion was induced for 24 hrs prior to fixation, expansion and imaging. BCC4 degradation leads to formation of double head parasites due to a defective BC. MORN1 is still able to localize to the basal end of daughter cells but ring integrity is lost after BCC4 degradation. Magnified panels 1 and 2 correspond with the boxed areas in the left and right panel with the corresponding number. BC, basal complex; cc, centrocone (spindle pole). 3D rendered rotations of these parasites are provided in Supplementary Movies S4 and S5. **f.** Exogenous MORN1-mCherryRFP overexpression driven by the strong constitutive α-tubulin promoter accumulates MORN1 in the basal complex and disrupts IMC formation but leaves subpellicular microtubule cytoskeleton formation intact (Gubbels et al., 2006). SR-SIM microscopy of parasites harboring endogenously tagged BCC4-3xMyc were transiently transfected with MORN1-mCherryRFP, fixed after overnight growth and stained with Myc andβ-tubulin antiserum. BCC4 does not accumulate with MORN1-mCherryRFP but becomes scattered largely along the subpellicular microtubules.

To dynamically track the BC upon BCC4 depletion, we performed time-lapse microscopy of BCC4-mAID parasites co-expressing endogenously YFP-tagged MORN1 (Fig 7d, Supplementary Movies 2 and 3). Parasites grown in absence of IAA exhibited BC formation and cell division dynamics as previously reported (Gubbels et al., 2006; Hu, 2008). We started imaging vacuoles containing two or four parasites in division when nascent MORN1 rings were visibly emerging around the MORN1 centrocone signal, early in S-phase. MORN1 signal at the apical end of nascent daughter cells was detected approximately 24 to 30 min after imaging started and communicated the basal to apical expansion of daughters (Fig 7d open arrow heads; Supplementary Movie S2). From here, the BC initially expanded until approximately minute 54 and then moves with the basal end of forming daughters, surpassing the centrocone between 60 and 66 mins (Fig 7d closed arrowheads). Contrary to these dynamics, parasites that were treated with IAA 2 hrs before imaging failed to produce visible daughter bud apical ends by minute 45. The daughter BCs formed close to the centrocones as seen in controls and showed an initial expansion (Fig. 7d blue arrowheads). However, at 54 mins two of the four BCs started to fragment, and by 75 min all BCs were fragmented (red arrowheads). In summary, in absence of BCC4, MORN1 is initially assembled into the BC, but midway through daughter bud assembly the MORN1 signal fragments and the BC falls apart. The critical loss of MORN1 at this point phenocopies, if not exaggerates, the MORN1-depleted phenotype and prevents further tapering of the daughters during the division process (Heaslip et al., 2010; Lorestani et al., 2010).

To gain higher resolution information of the BC fragmentation phenotype and investigate how the BC interfaces with the microtubular cytoskeleton we applied ultrastructural expansion microscopy (U-ExM) as recently established for *T. gondii* (Tosetti et al., 2020). BCC4-mAID3xMyc parasites co-expressing YFP-MORN1 were co-stained with β-tubulin antiserum. Upon 24 hrs IAA induction we observed the bell-shaped daughter buds as seen with IMC staining (Fig. 7e vs. Fig. 6e, f), thereby implying that IMC and sub-pellicular microtubules remain associated. Although MORN1 still localizes to the (wider) basal end of the sub-pellicular microtubules, it lost its smooth and continuous appearance as seen for the undisturbed BC (Fig. 7e).

The continued association of MORN1 with the (+)-ends of daughter microtubules suggest a possible mechanism for BC positioning at the end daughter buds. Another way to interfere with daughter budding and BC positioning is through the overexpression of MORN1 itself (Gubbels et al., 2006). This induces the formation of MORN1 rings that are no longer anchored to the IMC, which as a result has an unstructured appearance. The formation of the sub-pellicular microtube cytoskeleton is unaffected, though the microtubule ends are not sequestered and are fraying, which results in severely misshapen parasite (Gubbels et al., 2006). When we transiently overexpressed mCherryRFP-MORN1 in BCC4 3xMyc parasites, MORN1 accumulated in three distinct spots corresponding with one mother and two daughter BCs (Fig. 7f). Under these conditions, BCC4 did not colocalize with MORN1 and failed to assemble in rings. In contrast, the BCC4 signal was dispersed over the entire parasite and foci appeared to be present along the sub-pellicular microtubules (Fig. 7f).

In conclusion, our data support a critical role for BCC4 together with MORN1 in the BC that need each other to stabilize the BC at the time MyoJ and Cen2 recruitment at the mid-budding point. However, in contrast to MORN1, BCC4 is only temporally needed till division is completed, upon which it is removed from the BC. These data highlight a critical new protein and event in the life cycle of the BC.

### BCC1 is essential for final tapering and recruits of MyoJ and Cen2 to the BC

Based on the primary structure, which only harbors several coiled-coil domains as identifiable motifs, function cannot be predicted for BCC1 (Fig. 8a). The association of BCC1 with MyoJ and Cen2 in BCSC-4 (Fig. 2b) suggested a potential role in the constriction and basal tapering step of the BC. Indeed, BCC1 tagged at the C-terminus with 3xMyc-tag is only recruited to the BC in the middle of budding, which mimics the dynamic of MyoJ and Cen2 recruitment (Fig. 8b). Furthermore, BCC1 co-localizes robustly with MyoJ and Cen2 and not with MORN1 (Fig. 8c). Thus, together with MyoJ and Cen2, BCC1 inhabits the most extreme basal BC area making up the posterior cup in non-dividing parasites.

**Fig 8.**
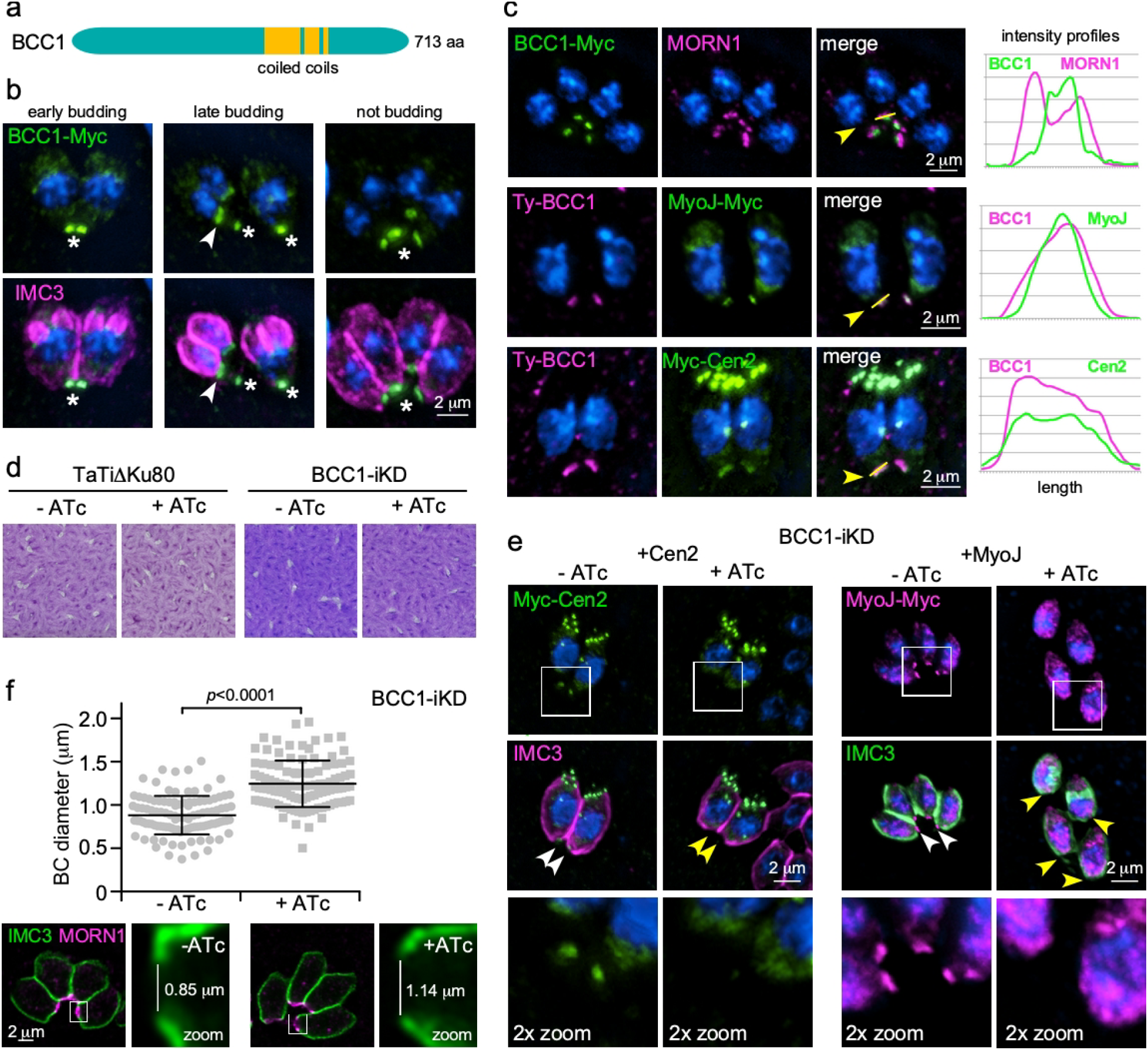
BCC1 acts in the final constriction phase of the BC. **a.** Schematic representation of BCC1, which only harbors coiled-coils as recognizable domains. **b.** BCC1-3xMyc associates with the BC in the second half of division, when daughter cells begin to taper. Asterisks mark BCC1 in the BC of the mature cytoskeleton; arrowheads mark BCC1 in the BC of the budding daughter cytoskeletons. DNA is stained with DAPI (blue). **c.** Colocalization of BCC1-3xMyc with TY-MORN1or TetO7sag4-TyBCC1with MyoJ-3xMyc or 2xMyc-Cen2 demonstrated that BCC1 resides in the most basal BC compartment defined by MyoJ and Cen2. Intensity profiles depict fluorescence intensity (y-axis) over length (yellow bar in adjacent image) on the x-axis. DNA is stained with DAPI (blue). **d.** Plaque assays of BCC1 knock-down by tetracycline-regulatable promoter (BCC1-iKD) show no significant reduction in viability over 7 days. **e.** BCC1 knockdown results in the loss of Cen2 and MyoJ from the mature BC. Yellow arrowheads point at the yellow line that corresponds with the intensity profiles on the right. Boxed regions in top panels are magnified in the lower panels. White arrowheads mark Cen2 or MyoJ in the basal complex; yellow arrowheads mark their absence. DNA is stained with DAPI (blue). **f.** BCC1 knockdown leads to an impaired constriction of the BC in the conclusion of cell division: BC diameter was measured as illustrated at the bottom panels. Boxed regions in left panels are magnified in the right panels. At least 100 mature BCs were measured and are plotted. Horizontal line marks the average; error bars represent sem.

BCC1 has a borderline lytic cycle fitness score of -1.88, suggesting it may or may not be an essential protein (Sidik et al., 2016). To test this directly we replaced the BCC1 promoter with the tetracycline-regulatable promoter and simultaneously inserted an N-terminal Ty-tag. Parasites grown for 7 days ±ATc showed no difference in plaque size, indicating that BCC1 is not essential for *in vitro* proliferation (Fig. 8d), again mirroring the phenotypes of MyoJ and Cen2 depletion (Frenal et al., 2017). Moreover, BCC1 depletion did prevent the association of both Cen2 and MyoJ with the BC, but Cen2 localization to the centrosome, apical end, and apical annuli was not affected (Fig. 8e). Morphologically, BCC1 depleted parasites appeared ‘stumpy’ due to loss of basal tapering. This phenotype is similar to depletion of MyoJ, Cen2 or actin, which are all defective in final BC constriction and likely the formation of the posterior cup (Heaslip et al., 2010; Frenal et al., 2017; Periz et al., 2017). Consistent with this model, we measured a 30% wider gap of the mature IMC’s basal end in BCC1-depleted parasites with an average basal diameter of 1.26±0.27 μm in BCC1-knockdown parasites compared to 0.89±0.22 μm in untreated control parasites (Fig. 8f). In conclusion, BCC1 is a novel protein required for recruiting MyoJ and Cen2 at the BC and is thereby essential for the final constriction of the BC in the conclusion of cell division.

## Discussion

The multi-bait proximity biotinylation approach paired with advanced statistical analysis provided a highly refined architectural map of the BC that we used as road map for guided functional dissection of the BC’s essential role in cell division. We successfully identified the two proteins (MORN1 and BCC4) that are the key controllers of BC stability throughout the cell division process. This led to a model wherein a motor protein is not essential, and the key maintaining ultrastructural integrity of a stretchable BC. In addition, we identified a new player in the MyoJ/Cen2 cluster (BCC1) that is essential to their recruitment to the BC in the second half of daughter division that supports the non-essential, final constriction step. Furthermore, our work uncovered several additional new insights pertaining to the hierarchical steps and structures in the unique cell division process. We demonstrate that the earliest structures of the daughter cytoskeleton are deposited in 5-fold symmetrical foci on which the apical cap assembles in apical direction, and the rest of the cytoskeleton including the BC assembles in the basal direction. These five foci define from the outset of budding the symmetry of the daughter buds, which contain 5 apical annuli at the suture between the cap-alveolus and 5 alveolar vesicle rows extending in basal direction. Moreover, we observed that the dynamic of sutures is interwoven in the assembly process, where BCC0 transitions and tracks with the assembling sutures. In addition, BCC3 tracks with the sutures as well and its later transition to the BC suggest this could provide a signaling step to maturation. Notably, both BCC0 and BCC3 are released from the BC upon cell division and therefore present a suture dynamic not reported before. In Fig. 9 we assembled a model that summarizes the new insights from this study.

**Fig 9.**
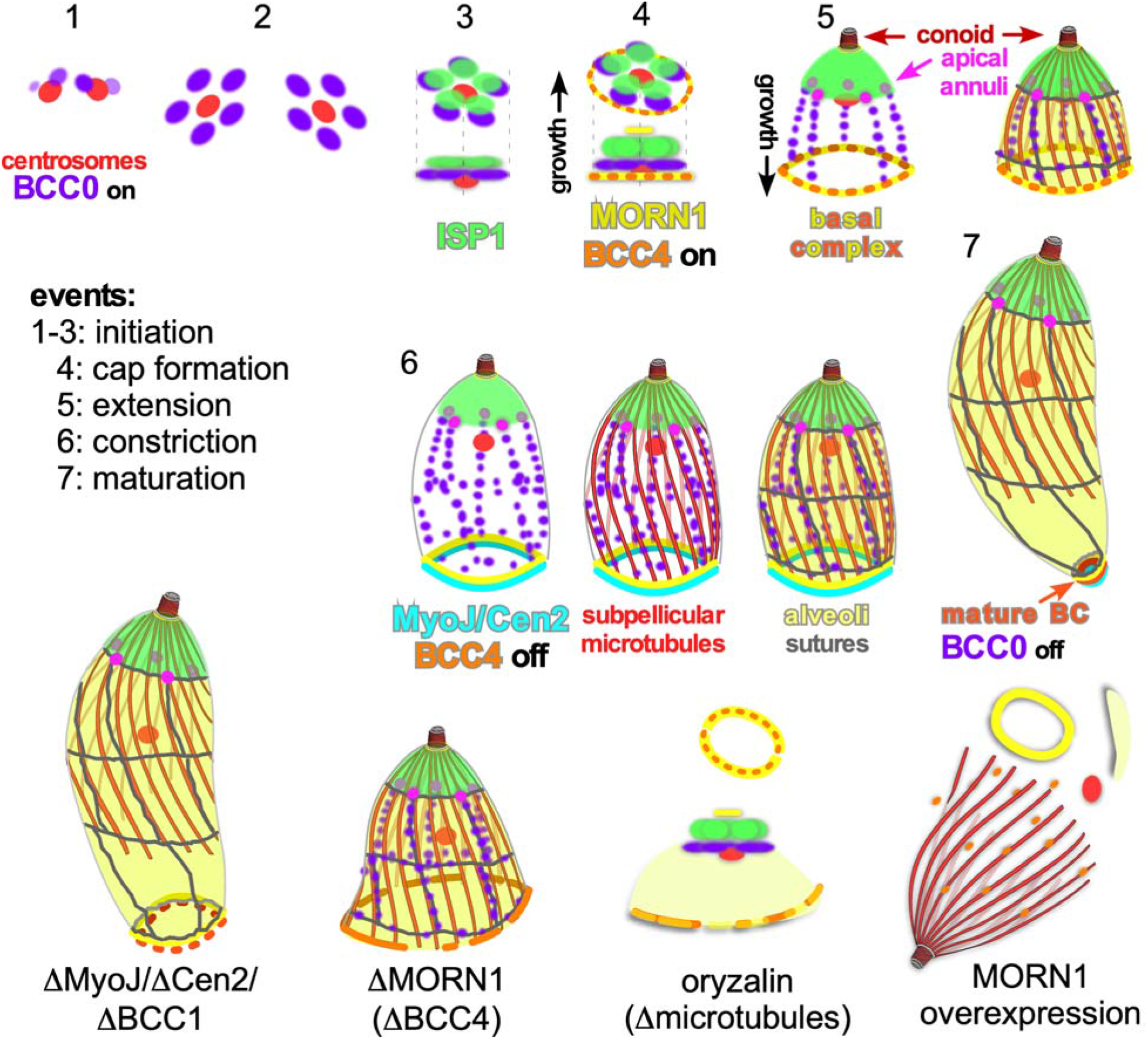
Summarizing schematic. Only defining components in BC formation and daughter budding are depicted. Note that ‘cap extension’ (step 4) and ‘extension’ (step 5) occur simultaneously but progress in opposite directions. The appearance of the name of components coincides with their recruitment to the BC, whereas for BC components that are dynamic, their named recruitment is defined by ‘on’ and their release by ‘off’. The schematics representing the phenotype mechanisms of various deletion mutants, chemical disruption (oryzalin), or overexpression (MORN1) are interpreted from the experimental data.

Regarding architecture of the BC, we firmly defined four different BCSCs which fit with the experimental co-localization experiments of defining BCSC components (Fig 1a; (Anderson-White et al., 2011)). We can assign the most critical function to BCSC-containing MORN1 and BCC4, as they are essential to maintain BC integrity which is of vital importance to successfully complete cell division. There are no apparent specific or dedicated functions organized in BCSC2 or 3 as they do not contain any BC localizing proteins with fitness scores hinting at essential roles, other than that they are possibly critical to interface the BC with IMC cytoskeleton and therefore might provide a complex level of support (Fig. 2, S2). BCSC-4 on the other hand harbors three key proteins (BCC1/MyoJ/Cen2) needed to taper the parasites in the finalizing steps of cell division.

Although temporal resolution could not be gleaned from the statistical analysis of the BioID data, we addressed this by tagging genes and tracking their localization throughout daughter development. The collective insights are presented in Fig. S3, which highlight four distinct protein recruitment steps to the BC coinciding with the functional steps in the BC: initiation, extension, constriction, and maturation. Although initiation exclusively contains BCSC-1 proteins, the other three steps sample from all four BCSCs, supporting a parallel BCSC development model, rather than a sequential model. Although the majority of proteins remains associated with the BC upon maturation, several early recruited proteins are released from the BC, which support a role unique to cell division. To date, a potential function for the proteins recruited to the BC in the mature cytoskeleton has been elusive, other than this could potentially be related to differentially control the stability of mother and daughter parasite during the division process (Gubbels et al., 2021).

Regarding the initiation of daughter budding, daughter scaffold proteins are recruited to the outer core of the duplicated centrosomes (Suvorova et al., 2015). Several known proteins recruited very early are IMC32 (Torres et al., 2021), AC9 (O’Shaughnessy et al., 2020; Tosetti et al., 2020), and FBXO1 (Baptista et al., 2019), for both of which a 5-fold symmetry can be appreciated as we observed for BCC0 and BCC3. IMC32 and FBXO1 are required for IMC membrane skeleton formation, where AC9 is critical conoid and sub-pellicular microtubule formation. Here we show that BCC0 and BCC3 also seem to define the position of the sutures and most likely the apical annuli, and thus are pivotal for defining the correct alveolar vesicle architecture. Moreover, the detection of BCC0 in our BC-BioID screen is contrasted against the absence of IMC32, AC9 and FBXO1 from this data set, which strongly supports that BCC0 is foundational for the BC. Further catering to this assessment is that BCC0 has been posited as a potential ortholog as a putative radial spoke protein 5 (RSP5) ortholog of RSP5 in *C. reinhardtii*) which binds to MORN domain proteins, though the statistical support was moderate at best (Portman et al., 2014; Portman and Slapeta, 2014).

One of the most striking observation is that bud growth is bi-directional from the foundation laid down by BCC0 and BCC3: ISP1 representing the apical cap assembles and extends in the apical direction (Fig. 3c, Fig. 9, panels 4 and 5), whereas the basal complex marked by MORN1, the longitudinal sutures marked by both BCC0 and 3 as well as the more basal alveoli marked by IMC3 extend in the basal direction (Fig. 3c, d, Fig. 4b-d, Fig. 9, panels 4 and 5). This is counter to the going model where extension is considered to occur exclusively in the basal direction (Anderson-White et al., 2012; Francia and Striepen, 2014). However, our revised model fits with an observatios reported that did not immediately align with the old model: upon ISP2 depletion, ISP1 still assembles in rings around the centrosome, but budding does not proceed any further (Beck et al., 2010). Our interpretation is that ISP1 gets stuck on the 5-fold symmetrical scaffold, but ISP2 is needed to extend the cytoskeleton either in apical or basal direction, or both. Furthermore, the role of the orthologous ISP proteins gleaned from *Plasmodium* ookinete formation, which pushes cytoskeleton buds outward from the plasma membrane, has provided a model of a protein palmitoylation driven outward push to generate the cup shape, which can then extend (Wang et al., 2020). We think this outward push principle likely also applies in *T. gondii*, but only to the apical cap formation in apical direction. Further support is that both the apical cap and the ookinete membrane skeleton are composed of only a single alveolar vesicle. The more complex alveolar and suture quilt seen in *T. gondii* seems to rely on a different assembly principle, notably the addition of components from the basal end of the bud (Ouologuem and Roos, 2014). Furthermore, DHHC14, which we detected in BCSC-2, which also contains suture proteins and IMC components (Fig. S2) is critical to daughter cytoskeleton assembly (Dogga and Frenal, 2020) and fits with our model. Thus, we propose a novel bi-directional model of *T. gondii* tachyzoite assembly, where the BC is the major factor in basal direction daughter bud growth.

The other major novel insight regarding the critical function of the BC in daughter cell assembly is that its most critical function is to keep the basal end of the daughter bud together. This becomes critical at the midpoint of daughter budding when the nucleus is entering the daughter buds. At this same time, many different proteins are recruited to the BC (Fig. S3), including the BCC1/MyoJ/Cen2 complex. We considered that the BC might be critical to keep the (+)-ends of the microtubules together, e.g. mimicking the MORN1 overexpression phenotype where in absence of the BC (and IMC), the sub-pellicular microtubules appear like an opened umbrella (Fig. 7f; (Gubbels et al., 2006)). This would imply that during the first half of division the BC is permitted to expand like a rubber band, but that the band reaches its maximal stretch at the midpoint of cell division, upon which it needs to be reinforced with additional proteins. This model explains the fragmentation of the BC upon BCC4 or MORN1 deletion leading to the double-headed phenotype (Fig. 7a-e). The recent report that MORN1 dimers can attain two different confirmations, either exhibiting an extended or a V-shape conformation, provides further support for a stretchable MORN1 ring (Sajko et al., 2020).

Beyond this step an additional transition in the daughter bud can be appreciated. When the parasite reaches about 2/3 of its final length the sub-pellicular microtubules stop extending. By U-ExM we actually observe that not all microtubules stop expanding at the same time (Fig. 7e and reported in (Gubbels et al., 2021)). We have been able to visualize this event as a very brief and transient appearance of (+)-end microtubule binding protein EB1 at the basal complex at this time (Fig. S3; (Chen et al., 2015b)). In our working model this is likely related to the final constriction by MyoJ/Cen2.

Another important implication of the BC-stretch model is that a motor protein is not needed, and indeed we did not find any support for any known contractile protein at the BC before the recruitment of BCC1/MyoJ/Cen2, whose depletion causes a vastly different and largely non-lethal phenotype (Fig. 8d-f). Notably, the actinomyosin ring in *Toxoplasma* tachyzoite cell division is not really required, and apicomplexans like *Plasmodium falciparum* seem to have done away with it (Rudlaff et al., 2019; Morano and Dvorin, 2021). Indeed, many protozoa complete cell division without an actinomyosin ring (Balasubramanian et al., 2012; Hammarton, 2019). We hypothesize that the actinomyosin ring together with Cen2 has evolutionary shared ancestry with “transition zone” at the base of motile cilia, sensory cilia and the connecting cilium in photoreceptor cells (Trojan et al., 2008). Particularly in photoreceptor cells, centrin/G-protein complexes organize signaling in retinal photoreceptor cells, next to a function as a barrier and transport of vesicles (Trojan et al., 2008). In Apicomplexa this function may still be intact (e.g. in recycling material from the residual body), and appears to have acquired an additional role in completing the last step of cell division.

Studies on the BC in *P. falciparum* have identified as set of proteins that is largely not conserved in *T. gondii* (Rudlaff et al., 2019). A notable player is PfCINCH, a dynamin-like protein with an essential role in *P. falciparum* BC constriction. In *T. gondii* DrpC is present in the BC but is involved in vesicular transport and partitioning of the mitochondrion late in cell division rather than BC constriction (Heredero-Bermejo et al., 2019; Melatti et al., 2019). We do detect DrpB in our data set (Fig. 2a) but this is likely a spurious hit as DrpB is involved in apical secretory organelle biogenesis (Breinich et al., 2009). However, BCC10 displays some homology with atlastin, which in vertebrates is a dynamin-like GTPase required for homotypic fusion of endoplasmic membrane tubules (Betancourt-Solis et al., 2018). However, we do not detect any orthology of BCC10 in *P. falciparum* (Fig. S4) whereas the mild fitness score indicates BCC10 is not essential, suggesting this not a functional PfCINCH ortholog.

To obtain a bird’s eye view of the conservation of the new BCCs we searched for putative orthologs in representative Apicomplexa by BLAST searched EuPathDB (Warrenfeltz et al., 2018). Interestingly, BCC conservation partitioned into three distinct groups (Fig. S4): group 1 is widely conserved across Apicomplexa, whereas group 2 is conserved across all Coccidia and the group 3 is narrowly conserved in tissue cyst forming Coccidia dividing by internal budding. Although group 1 contains a kinase (BCC2) and a phosphatase (BCC5) where the catalytic domain likely provides conservation, BCC9 lacks any identifiable domain and thus seems under strong selective pressure, yet has a fitness score of -1.36 suggesting it is not essential. The proteins in group 2 have mostly some identifiable motifs, but the proteins in group 3 are all hypothetical. Their lack of conserved motifs, high incidence of low complexity and coiled-coil regions, together with relatively low density of introns indicates these proteins are fast evolving and therefore identifying functional orthologs based on primary sequence in other species is very challenging, but importantly does not exclude the presence of functional orthologs. An alternative argument can be made that the lack of pan-apicomplexan conservation illustrates the differences in division modes (Gubbels et al., 2020). And although group 1 and 3 partition along the principally different internal versus external budding schism, group 2 bridges between these two division modes, suggesting it is at an evolutionary cross-road.

Taken together, our work has highlighted several tantalizing new insights in the hierarchical steps and structure of daughter cell assembly together with revealing a rubber-band model as mechanism of the BC’s essential role during cell division.

## Material and Methods

### Parasites and mammalian cell lines

Transgenic derivatives of the RH strain were maintained and assessed in hTERT immortalized human foreskin fibroblasts (HFF) except for IFA assays, which were performed in primary HFF cells, largely as previously described (Roos et al., 1994). Stable parasite transfectants were selected under 1 μM pyrimethamine, 20 µM chloramphenicol, 20 µM 5’-fluo-2’deoxyuridine (FUDR), or a combination of 25 mg/ml mycophenolic acid and 50 mg/ml xanthine (MPA/X).

### Plasmids and parasite strain generation

For endogenous 5’-end tagging of BC genes with BioID2 we used a previously reported method (Engelberg et al., 2020). In short, expression of the selection marker was linked to the integration into a specific gene locus (selection-linked integration (SLI) (Birnbaum et al., 2017)). A PCR amplicon consisting of the HXGPRT ORF linked to the sequence of the ty-1-bioid2 ORF by the T2A skip peptide sequence was transfected together with a CRISPR/Cas9 plasmid that generated a specific DNA double strand break around the ATG of the respective gene locus. The PCR amplicon carries 35 bp homologous flanks on each site to facilitate homologues repair. Transfected parasites were selected using MPA/X for expression of HXGPRT under the respective endogenous promoter.

Prey genes were analyzed by endogenous tagging via 3’-end replacement. Homologous 3’end flanks of a given prey gene were cloned via PmeI and AvrII into the integration plasmid to generate 3xMyc or YFP tagged alleles. 50 µg of plasmid DNA was linearized with a restriction enzyme digest before transfection in RHΔKu80 parasites.

Conditional knockdown parasite lines were generated by cloning a homologous 5’-flank of the respective ORF via BglII and NotI restriction sites into a DHFR-TetO7-sag4-Ty plasmid. Plasmid was linearized using a single restriction enzyme cut site and subsequently transfected in TATiΔKu80 parasites (Sheiner et al., 2011).

Endogenous 3’end tagging with mAID were done as follows. The DNA sequence of the minimal Auxin inducible degron (mAID) (Kubota et al., 2013) was amplified from a donor plasmid (kind gift of Dr. Lourido, Whitehead Institute) and cloned in frame with the PmeI/ArvII restriction sites of the 3xMyc-DHFR 3’end integration plasmid. The homologous 3’flank used for endogenous tagging of TGGT1_229260 with 3xMyc was cloned into the newly established plasmid and linearized as previously mentioned 50 µg of plasmid DNA was used for transfection of RH Tir1-3xFLAG parasites (Brown et al., 2017) and transfected cell lines were selected with Pyrimethamine. To induce specific protein degradation, parasites were incubated with 500 µM IAA (in 100% ethanol) for times indicated in the result section. To endogenously tag MORN1 or BCC4 in RH Tir1-3xFLAG parasites we generated a plasmid that linked the HXGPRT or DHFR/TS ORF via the T2A skip peptide to the YFP ORF, the 3xMyc or 5xV5 epitope tag sequence. A PCR amplicon was transfected together with 40 µg of a CRISPR/Cas9 plasmid that generated a DNA double strand break around the ATG of either MORN1 or BCC4. Parasites expressing YFP-MORN1 or 3xMyc-BCC4 were selected with MPA/Xan or Pyr as mentioned above.

### BioID sample preparation and mass spectrum analysis

Biotin labeling was done in two biological replicates (+biotin condition) and one biological replicate (-biotin condition) as reported before (Engelberg et al., 2020). Each biological replicate was run as two technical replicates on the mass spectrometer. Biotinylated proteins were identified as previously reported (Engelberg et al., 2020). In short, parasites expressing BioID2-fusion proteins were grown overnight ±150 μM biotin and extracellular parasites were harvested, filtered through a 3 mm membrane and mechanical lysed in 1%SDS in resuspension buffer (150 mM NaCl, 50 mM Tris-HCl pH 7.4). 1.5 mg of total protein lysate was used for streptavidin pull-down with Streptavidin-agarose beads (Fisher). Proteins bond on beads were reduced, alkylated and digested with Trypsin. Digested peptides were used for LC-MS/MS analysis, which was performed on an LTQ-Orbitrap Discovery mass spectrometer (ThermoFisher) coupled to an Agilent 1200 series HPLC.

The tandem MS data were searched using the SEQUEST algorithm (Eng et al., 1994) using a concatenated target/decoy variant of the *Toxoplasma gondii* GT1 ToxoDB-V29 database. A static modification of +57.02146 on cysteine was specified to account for alkylation by iodoacetamide. SEQUEST output files were filtered using DTASelect 2.0 (Tabb et al., 2002). Reported peptides were required to be unique to the assigned protein (minimum of two unique peptides per protein) and discriminant analyses were performed to achieve a peptide false-positive rate below 5%. Data for Ty-BioID2-Cen2 and YFP-BioID2 were used from a previous study (Engelberg et al., 2020).

### Analysis of mass-spec data by probabilistic calculation of interactions

Spectral counts of unique proteins were used to determine probability of interaction for given bait and preys using SAINTexpress (Teo et al., 2014). SAINTexpress was executed using the – L4 argument, compressing the four largest quantitative control values of a given prey in one virtual control. The resulting SAINTexpress matrix was visualized using the Prohits-viz online suite (Knight et al., 2017) with the following settings: abundance column set to “AverageSpec”, score column set to “false discovery rate (FDR)”, score filter = 0.1, secondary score filter = 0.25, lower abundance cutoff for prey correlation = “10 spectral counts” to generate the prey-prey correlation in Fig. 2a. The following preys were manually deleted from the analysis using the Zoom function in Prohits-viz: HXGPRT, TGGT1_269600 (annotated as biotin enzyme), TGGT1_289760 (annotated as biotin-synthase). The dot plot was generated by including a cytosolic BioID2-YFP control (Engelberg et al., 2020) in the SAINT analysis and setting the abundance column to “average Spectral Counts”, the score filter to “false discovery rate (FDR)”, the primary filter = “0.01”, the secondary filter = “0.05” and the log transformation = “log2”. Preys were manually arranged into BC sub complexes for Fig. 2b, the entire resulting dot plot is shown in Fig. S2b.

### Control-subtracted length-adjusted spectral counts

Normalized spectral counts (NormSpC) were calculated as previously reported (Youn et al., 2018; Go et al., 2021) using the following formula: 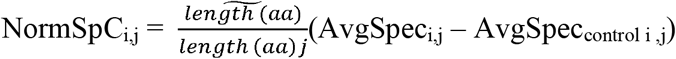. This resulted in control-subtracted length-adjusted spectral counts. For a given bait *i* and prey *j*, we first subtracted the average spectral counts for *j* detected within bait *i* (“no biotin” samples) from the average spectral counts detected for *j* with bait *i* when biotin was added. We then multiplied the difference with the quotient of the median length of all preys detected in bait *i* divided by the length of *j*. The heatmap visualizing NormSpC was generated in R using the ComplexHeatmap package (Gu et al., 2016).

### (Immuno-) fluorescence microscopy

Indirect immunofluorescence assays were performed on intracellular parasites grown overnight in 6-well plate containing coverslips confluent with HFF cells fixed with 100% methanol (unless stated otherwise) using the following primary antisera: mouse α-Ty clone BB2 (1:500; kindly provided by Dr. Lourido, Whitehead Institute), MAb 9E10 α-cMyc (1:50; Santa Cruz Biotechnology), MAb 9B11 α-cMyc Alexa488 (A488) conjugated (1:100; Cell Signaling Technologies), mouse α-V5 clone SV5-Pk1 (1:500, BioRad), rabbit α-®-tubulin (1:1,000; kindly provided by Naomi Morrissette, University of California, Irvine; (Morrissette and Sibley, 2002)), rat α-IMC3 (1:2,000; (Anderson-White et al., 2011)), rabbit α-IMC3 (1:2,000; generated against the N-terminal 120 amino acids fused to His6 generated as described in (Anderson-White et al., 2011)), rabbit α-human-Centrin2 (1:1,000; kindly provided by Iain Cheeseman, Whitehead Institute), and guinea pig α-AAP4 (1:200) (Engelberg et al., 2020). Streptavidin-A594 (1:1000; Thermo Fisher), A488 or A594 conjugated goat α-mouse, α-rabbit, α-rat, or α-guinea pig were used as secondary antibodies (1:500; Invitrogen). DNA was stained with 4’,6-diamidino-2-phenylindole (DAPI). A Zeiss Axiovert 200 M wide-field fluorescence microscope was used to collect images, which were deconvolved and adjusted for phase contrast using Volocity software (Improvision/Perkin Elmer). SR-SIM or Zeiss Airyscan imaging was performed on intracellular parasites fixed with 4% PFA in PBS and permeabilized with 0.25% TX-100 or fixed with 100% methanol. Images were acquired with a Zeiss ELYRA S.1 (SR-SIM) and Airyscan system in the Boston College Imaging Core in consultation with Bret Judson. All images were acquired, analyzed and adjusted using ZEN software and standard settings. Final image analyses were made with FIJI software.

### Live cell microscopy

Live cell microscopy of RH Tir1-3xFLAG BCC4mAID-Myc parasites expressing YFP-MORN1 was done using a Zeiss LSM880 with Airyscan unit in the Boston College Imaging Core in consultation with Bret Judson. Parasite dynamics were recorded using the “Airyscan fast” settings, in an incubation chamber set to 37°C. Parasites were grown overnight under standard culture conditions in 3 ml live cell dishes (MatTek). On the next day, culture medium was replaced with live cell imaging medium (DMEM without phenol red, 20 mM HEPES pH 7.4, 1% FBS, Penicillin/Streptomycin and Fungizone). To induce protein degradation, parasites were treated with 500 µM IAA (in 100% ethanol) 2 hrs before imaging started. The resulting data were deconvolved using standard Airyscan settings and movies processed with FIJI software.

### Quantification of daughter basal end diameter

BCC4-mAID-3xMyc parasites endogenously expressing YFP-MORN1, Ty-mAID-MORN1 parasites endogenously expressing 3xMyc-BCC4 or TirI parental parasites were seeded on coverslips with HFF host cells and grown over night. At the next day, protein degradation was initiated with 500 nM IAA and parasites were incubated for two more hours. Cells were fixed with 4% PFA and stained with anti-IMC3 and anti-Myc (in case of Ty-mAID-MORN1 parasites) serum and images of dividing parasites were acquired using the Zeiss LSM880 Airyscan. Images were analyzed in FIJI (Schindelin et al., 2012) and basal diameter for at least 50 daughter basal ends measured. Resulting data were visualized with the ggplot package in R (Wickham, 2016).

### Expansion Microscopy

Expansion of *Toxoplasma* tachyzoites was achieved by following recently published protocols (Gambarotto et al., 2019; Le Guennec et al., 2020; Tosetti et al., 2020). Briefly, tachyzoites growing in HFF cells for 24 hrs ±IAA were released by syringe lysis and filtered through a 12 μm membrane. Free tachyzoites were allowed to settle on poly-L-lysine (coated coverslips at 4°C for 30 min, followed by fixation in −20°C methanol for 7 min. The U-ExM protocol was started by incubating cover slips in 2xsolution (2% formaldehyde, 1.4% acrylamide (AA) in PBS) for five hours at 37°C. Gelation was done in Monomere solution (19% (w/w) sodium acrylate, 10% (w/w) AA and 0.1% (w/w) BIS-AA in PBS) complemented with ammonium persulfate (APS) and tetramethylethylenediamine (TEMED) for 1 hrs at 37°C, followed by incubation in denaturation buffer (200 mM SDS, 200 mM NaCl, 50 mM Tris, pH 9) at 95°C for 90 mins. Gels were incubated for a first round of expansion in ddH_2_O overnight and washed twice in PBS on the next day. As a primary antibody, mouse α beta-tubulin (12G10) and rabbit α-GFP (Torrey Pines) were used to stain parasite microtubules and YFP-MORN1 respectively. Gels were incubated in 2% BSA in PBS with primary antibodies at 37°C for three hours, washed three times with PBST (1xPBS + 0.1% Tween20) and incubated for three hours at 37°C in 2% BSA in PBS complemented with secondary antibody (goat anti-rabbit-Oregon Green, goat anti-mouse-A594, Invitrogen). Gels were washed three times in PBST before a second overnight expansion in ddH_2_O. For imaging, gels were mounted in 35 mm glass bottom microwell dishes (MatTek) and imaged on a Zeiss LSM880 with Airyscan unit using standard settings for image acquisition and Airyscan deconvolution. All imaging was done in the Boston College Imaging Core with help/advice of Dr. Bret Judson.

## Supporting information

Supplementary matrrial

## Acknowledgements

We thank Sudeshna Saha, Ciara Bauwens, and Adam Mehal for technical support, Bret Judson and the Boston College Imaging Core for infrastructure and support and fruitful discussions, Drs. Cheeseman, Lourido, and Morrissette for sharing reagents.

This study was supported by National Science Foundation (NSF) Major Research Instrumentation grant 1626072, National Institute of Health grants AI110690, AI128136, AI144856, and AI152387 a Deutsche Forschungsgemeinschaft grant, an American Heart Association post-doctoral fellowship grant 17POST33670577 and a Knights Templar Eye Foundation early career starter grant. The funders had no role in study design, data collection and analysis, decision to publish, or preparation of the manuscript.

## Author Contributions

KE and MJG conceived the approach, analyzed and interpreted the data, and co-wrote the manuscript. KE designed and generated all *T. gondii* mutants and data. TB and EW performed and analyzed the mass spectrometry data. CM assisted with data analysis, presentation and assessed the conservation of the BCCs. All authors proofread the manuscript.

## References

Anderson-White, B.R., Beck, J.R., Chen, C.T., Meissner, M., Bradley, P.J., and Gubbels, M.J. (2012). Cytoskeleton assembly in Toxoplasma gondii cell division. Int. Rev. Cell Mol. Biol. 298, 1–31.

Anderson-White, B.R., Ivey, F.D., Cheng, K., Szatanek, T., Lorestani, A., Beckers, C.J., Ferguson, D.J., Sahoo, N., and Gubbels, M.J. (2011). A family of intermediate filament-like proteins is sequentially assembled into the cytoskeleton of Toxoplasma gondii. Cell Microbiol 13, 18–31.

Balasubramanian, M.K., Srinivasan, R., Huang, Y., and Ng, K.H. (2012). Comparing contractile apparatus-driven cytokinesis mechanisms across kingdoms. Cytoskeleton 69, 942–956.

Bang, W., Kim, S., Ueda, A., Vikram, M., Yun, D., Bressan, R.A., Hasegawa, P.M., Bahk, J., and Koiwa, H. (2006). Arabidopsis carboxyl-terminal domain phosphatase-like isoforms share common catalytic and interaction domains but have distinct in planta functions. Plant Physiol 142, 586–594.

Baptista, C.G., Lis, A., Deng, B., Gas-Pascual, E., Dittmar, A., Sigurdson, W., West, C.M., and Blader, I.J. (2019). Toxoplasma F-box protein 1 is required for daughter cell scaffold function during parasite replication. PLoS Pathog 15, e1007946.

Barylyuk, K., Koreny, L., Ke, H., Butterworth, S., Crook, O.M., Lassadi, I., Gupta, V., Tromer, E., Mourier, T., Stevens, T.J., Breckels, L.M., Pain, A., Lilley, K.S., and Waller, R.F. (2020). A Comprehensive Subcellular Atlas of the Toxoplasma Proteome via hyperLOPIT Provides Spatial Context for Protein Functions. Cell Host Microbe 28, 752–766.

Beck, J.R., Rodriguez-Fernandez, I.A., Cruz De Leon, J., Huynh, M.H., Carruthers, V.B., Morrissette, N.S., and Bradley, P.J. (2010). A novel family of Toxoplasma IMC proteins displays a hierarchical organization and functions in coordinating parasite division. PLoS Pathog 6, e1001094.

Betancourt-Solis, M.A., Desai, T., and Mcnew, J.A. (2018). The atlastin membrane anchor forms an intramembrane hairpin that does not span the phospholipid bilayer. J Biol Chem 293, 18514–18524.

Birnbaum, J., Flemming, S., Reichard, N., Soares, A.B., Mesen-Ramirez, P., Jonscher, E., Bergmann, B., and Spielmann, T. (2017). A genetic system to study Plasmodium falciparum protein function. Nat Methods 14, 450–456.

Breinich, M.S., Ferguson, D.J., Foth, B.J., Van Dooren, G.G., Lebrun, M., Quon, D.V., Striepen, B., Bradley, P.J., Frischknecht, F., Carruthers, V.B., and Meissner, M. (2009). A Dynamin Is Required for the Biogenesis of Secretory Organelles in Toxoplasma gondii. Curr Biol 19, 277–286.

Brown, K.M., Long, S., and Sibley, L.D. (2017). Plasma Membrane Association by N-Acylation Governs PKG Function in Toxoplasma gondii. MBio 8.

Chen, A.L., Kim, E.W., Toh, J.Y., Vashisht, A.A., Rashoff, A.Q., Van, C., Huang, A.S., Moon, A.S., Bell, H.N., Bentolila, L.A., Wohlschlegel, J.A., and Bradley, P.J. (2015a). Novel components of the Toxoplasma inner membrane complex revealed by BioID. MBio 6, e02357–02314.

Chen, A.L., Moon, A.S., Bell, H.N., Huang, A.S., Vashisht, A.A., Toh, J.Y., Lin, A.H., Nadipuram, S.M., Kim, E.W., Choi, C.P., Wohlschlegel, J.A., and Bradley, P.J. (2017). Novel insights into the composition and function of the Toxoplasma IMC sutures. Cell Microbiol 19.

Chen, C.T., and Gubbels, M.J. (2013). The Toxoplasma gondii centrosome is the platform for internal daughter budding as revealed by a Nek1 kinase mutant. J Cell Sci 126, 3344–3355.

Chen, C.T., Kelly, M., Leon, J., Nwagbara, B., Ebbert, P., Ferguson, D.J., Lowery, L.A., Morrissette, N., and Gubbels, M.J. (2015b). Compartmentalized Toxoplasma EB1 bundles spindle microtubules to secure accurate chromosome segregation. Mol Biol Cell 26, 4562–4576.

Choi, H., Larsen, B., Lin, Z.Y., Breitkreutz, A., Mellacheruvu, D., Fermin, D., Qin, Z.S., Tyers, M., Gingras, A.C., and Nesvizhskii, A.I. (2011). SAINT: probabilistic scoring of affinity purification-mass spectrometry data. Nat Methods 8, 70–73.

Dogga, S.K., and Frenal, K. (2020). Two palmitoyl acyltransferases involved sequentially in the biogenesis of the inner membrane complex of Toxoplasma gondii. Cell Microbiol 22, e13212.

Eng, J.K., Mccormack, A.L., and Yates, J.R. (1994). An approach to correlate tandem mass spectral data of peptides with amino acid sequences in a protein database. J Am Soc Mass Spectrom 5, 976–989.

Engelberg, K., Chen, C.T., Bechtel, T., Sanchez Guzman, V., Drozda, A.A., Chavan, S., Weerapana, E., and Gubbels, M.J. (2020). The apical annuli of Toxoplasma gondii are composed of coiled-coil and signalling proteins embedded in the inner membrane complex sutures. Cell Microbiol 22, e13112.

Engelberg, K., Ivey, F.D., Lin, A., Kono, M., Lorestani, A., Faugno-Fusci, D., Gilberger, T.W., White, M., and Gubbels, M.-J. (2016). A MORN1-assocatiated HAD phosphatase in the basal complex is essential for Toxoplasma gondii daughter budding. Cell Microbiol 18, 1153–1171.

Francia, M.E., and Striepen, B. (2014). Cell division in apicomplexan parasites. Nature reviews. Microbiology 12, 125–136.

Frenal, K., Jacot, D., Hammoudi, P.M., Graindorge, A., Maco, B., and Soldati-Favre, D. (2017). Myosin-dependent cell-cell communication controls synchronicity of division in acute and chronic stages of Toxoplasma gondii. Nat Commun 8, 15710.

Gajria, B., Bahl, A., Brestelli, J., Dommer, J., Fischer, S., Gao, X., Heiges, M., Iodice, J., Kissinger, J.C., Mackey, A.J., Pinney, D.F., Roos, D.S., Stoeckert, C.J., Jr., Wang, H., and Brunk, B.P. (2008). ToxoDB: an integrated Toxoplasma gondii database resource. Nucleic Acids Res 36, D553–556.

Gambarotto, D., Zwettler, F.U., Le Guennec, M., Schmidt-Cernohorska, M., Fortun, D., Borgers, S., Heine, J., Schloetel, J.G., Reuss, M., Unser, M., Boyden, E.S., Sauer, M., Hamel, V., and Guichard, P. (2019). Imaging cellular ultrastructures using expansion microscopy (U-ExM). Nat Methods 16, 71–74.

Go, C.D., Knight, J.D.R., Rajasekharan, A., Rathod, B., Hesketh, G.G., Abe, K.T., Youn, J.Y., Samavarchi-Tehrani, P., Zhang, H., Zhu, L.Y., Popiel, E., Lambert, J.P., Coyaud, E., Cheung, S.W.T., Rajendran, D., Wong, C.J., Antonicka, H., Pelletier, L., Palazzo, A.F., Shoubridge, E.A., Raught, B., and Gingras, A.C. (2021). A proximity-dependent biotinylation map of a human cell. Nature 595, 120–124.

Goodenough, U., Roth, R., Kariyawasam, T., He, A., and Lee, J.H. (2018). Epiplasts: Membrane Skeletons and Epiplastin Proteins in Euglenids, Glaucophytes, Cryptophytes, Ciliates, Dinoflagellates, and Apicomplexans. MBio 9, e02020–02018.

Gu, Z., Eils, R., and Schlesner, M. (2016). Complex heatmaps reveal patterns and correlations in multidimensional genomic data. Bioinformatics 32, 2847–2849.

Gubbels, M.J., Coppens, I., Zarringhalam, K., Duraisingh, M.T., and Engelberg, K. (2021). The modular circuitry of apicomplexan cell division plasticity. Front Cell Infect Microbiol 11, 670049.

Gubbels, M.J., Keroack, C.D., Dangoudoubiyam, S., Worliczek, H.L., Paul, A.S., Bauwens, C., Elsworth, B., Engelberg, K., Howe, D.K., Coppens, I., and Duraisingh, M.T. (2020). Fussing About Fission: Defining Variety Among Mainstream and Exotic Apicomplexan Cell Division Modes. Front Cell Infect Microbiol 10, 269.

Gubbels, M.J., Vaishnava, S., Boot, N., Dubremetz, J.F., and Striepen, B. (2006). A MORN-repeat protein is a dynamic component of the Toxoplasma gondii cell division apparatus. J Cell Sci 119, 2236–2245.

Hammarton, T.C. (2019). Who Needs a Contractile Actomyosin Ring? The Plethora of Alternative Ways to Divide a Protozoan Parasite. Front Cell Infect Microbiol 9, 397.

Heaslip, A.T., Dzierszinski, F., Stein, B., and Hu, K. (2010). TgMORN1 Is a Key Organizer for the Basal Complex of Toxoplasma gondii. PLoS Pathog 6, e1000754.

Heredero-Bermejo, I., Varberg, J.M., Charvat, R., Jacobs, K., Garbuz, T., Sullivan, W.J., Jr., and Arrizabalaga, G. (2019). TgDrpC, an atypical dynamin-related protein in Toxoplasma gondii, is associated with vesicular transport factors and parasite division. Mol Microbiol 111, 46–64.

Hu, K. (2008). Organizational changes of the daughter basal complex during the parasite replication of Toxoplasma gondii. PLoS Pathog 4, e10.

Hu, K., Johnson, J., Florens, L., Fraunholz, M., Suravajjala, S., Dilullo, C., Yates, J., Roos, D.S., and Murray, J.M. (2006). Cytoskeletal components of an invasion machine--the apical complex of Toxoplasma gondii. PLoS Pathog 2, e13.

Knight, J.D.R., Choi, H., Gupta, G.D., Pelletier, L., Raught, B., Nesvizhskii, A.I., and Gingras, A.C. (2017). ProHits-viz: a suite of web tools for visualizing interaction proteomics data. Nat Methods 14, 645–646.

Kubota, T., Nishimura, K., Kanemaki, M.T., and Donaldson, A.D. (2013). The Elg1 replication factor C-like complex functions in PCNA unloading during DNA replication. Mol Cell 50, 273–280.

Le Guennec, M., Klena, N., Gambarotto, D., Laporte, M.H., Tassin, A.M., Van Den Hoek, H., Erdmann, P.S., Schaffer, M., Kovacik, L., Borgers, S., Goldie, K.N., Stahlberg, H., Bornens, M., Azimzadeh, J., Engel, B.D., Hamel, V., and Guichard, P. (2020). A helical inner scaffold provides a structural basis for centriole cohesion. Sci Adv 6, eaaz4137.

Lentini, G., Kong-Hap, M., El Hajj, H., Francia, M., Claudet, C., Striepen, B., Dubremetz, J.F., and Lebrun, M. (2015). Identification and characterization of Toxoplasma SIP, a conserved apicomplexan cytoskeleton protein involved in maintaining the shape, motility and virulence of the parasite. Cellular Microbiology 17, 62–78.

Leveque, M.F., Berry, L., and Besteiro, S. (2016). An evolutionarily conserved SSNA1/DIP13 homologue is a component of both basal and apical complexes of Toxoplasma gondii. Sci Rep 6, 27809.

Lorestani, A., Ivey, F.D., Thirugnanam, S., Busby, M.A., Marth, G.T., Cheeseman, I.M., and Gubbels, M.J. (2012). Targeted proteomic dissection of Toxoplasma cytoskeleton sub-compartments using MORN1. Cytoskeleton 69, 1069–1085.

Lorestani, A., Sheiner, L., Yang, K., Robertson, S.D., Sahoo, N., Brooks, C.F., Ferguson, D.J., Striepen, B., and Gubbels, M.J. (2010). A Toxoplasma MORN1 Null Mutant Undergoes Repeated Divisions but Is Defective in Basal Assembly, Apicoplast Division and Cytokinesis. PLoS ONE 5, e12302.

Mann, T., and Beckers, C. (2001). Characterization of the subpellicular network, a filamentous membrane skeletal component in the parasite Toxoplasma gondii. Mol Biochem Parasitol 115, 257–268.

Meissner, M., Schluter, D., and Soldati, D. (2002). Role of Toxoplasma gondii myosin A in powering parasite gliding and host cell invasion. Science 298, 837–840.

Melatti, C., Pieperhoff, M., Lemgruber, L., Pohl, E., Sheiner, L., and Meissner, M. (2019). A unique dynamin-related protein is essential for mitochondrial fission in Toxoplasma gondii. PLoS Pathog 15, e1007512.

Montoya, J.G., and Liesenfeld, O. (2004). Toxoplasmosis. Lancet 363, 1965–1976.

Morano, A.A., and Dvorin, J.D. (2021). The Ringleaders: Understanding the Apicomplexan Basal Complex Through Comparison to Established Contractile Ring Systems. Front Cell Infect Microbiol 11, 656976.

Morrissette, N.S., and Sibley, L.D. (2002). Disruption of microtubules uncouples budding and nuclear division in Toxoplasma gondii. J Cell Sci 115, 1017–1025.

Nishi, M., Hu, K., Murray, J.M., and Roos, D.S. (2008). Organellar dynamics during the cell cycle of Toxoplasma gondii. J Cell Sci 121, 1559–1568.

O’shaughnessy, W.J., Hu, X., Beraki, T., Mcdougal, M., and Reese, M.L. (2020). Loss of a conserved MAPK causes catastrophic failure in assembly of a specialized cilium-like structure in Toxoplasma gondii. Mol Biol Cell 31, 881–888.

Ouologuem, D.T., and Roos, D.S. (2014). Dynamics of the Toxoplasma gondii inner membrane complex. J Cell Sci 127, 3320–3330.

Periz, J., Whitelaw, J., Harding, C., Gras, S., Del Rosario Minina, M.I., Latorre-Barragan, F., Lemgruber, L., Reimer, M.A., Insall, R., Heaslip, A., and Meissner, M. (2017). Toxoplasma gondii F-actin forms an extensive filamentous network required for material exchange and parasite maturation. Elife 6.

Portman, N., Foster, C., Walker, G., and Slapeta, J. (2014). Evidence of intraflagellar transport and apical complex formation in a free-living relative of the apicomplexa. Eukaryot Cell 13, 10–20.

Portman, N., and Slapeta, J. (2014). The flagellar contribution to the apical complex: a new tool for the eukaryotic Swiss Army knife? Trends Parasitol 30, 58–64.

Roos, D.S., Donald, R.G., Morrissette, N.S., and Moulton, A.L. (1994). Molecular tools for genetic dissection of the protozoan parasite Toxoplasma gondii. Methods Cell Biol 45, 27–63.

Rudlaff, R.M., Kraemer, S., Streva, V.A., and Dvorin, J.D. (2019). An essential contractile ring protein controls cell division in Plasmodium falciparum. Nat Commun 10, 2181.

Sajko, S., Grishkovskaya, I., Kostan, J., Graewert, M., Setiawan, K., Trubestein, L., Niedermuller, K., Gehin, C., Sponga, A., Puchinger, M., Gavin, A.C., Leonard, T.A., Svergun, D.I., Smith, T.K., Morriswood, B., and Djinovic-Carugo, K. (2020). Structures of three MORN repeat proteins and a re-evaluation of the proposed lipid-binding properties of MORN repeats. PLoS One 15, e0242677.

Schindelin, J., Arganda-Carreras, I., Frise, E., Kaynig, V., Longair, M., Pietzsch, T., Preibisch, S., Rueden, C., Saalfeld, S., Schmid, B., Tinevez, J.Y., White, D.J., Hartenstein, V., Eliceiri, K., Tomancak, P., and Cardona, A. (2012). Fiji: an open-source platform for biological-image analysis. Nat Methods 9, 676–682.

Shaw, M.K., Compton, H.L., Roos, D.S., and Tilney, L.G. (2000). Microtubules, but not actin filaments, drive daughter cell budding and cell division in Toxoplasma gondii. J Cell Sci 113 ( Pt 7), 1241–1254.

Sheiner, L., Demerly, J.L., Poulsen, N., Beatty, W.L., Lucas, O., Behnke, M.S., White, M.W., and Striepen, B. (2011). A systematic screen to discover and analyze apicoplast proteins identifies a conserved and essential protein import factor. PLoS Pathog 7, e1002392.

Sidik, S.M., Huet, D., Ganesan, S.M., Huynh, M.H., Wang, T., Nasamu, A.S., Thiru, P., Saeij, J.P.J., Carruthers, V.B., Niles, J.C., and Lourido, S. (2016). A Genome-wide CRISPR Screen in Toxoplasma Identifies Essential Apicomplexan Genes. Cell 166, 1423–1435 e1412.

Skariah, S., Walwyn, O., Engelberg, K., Gubbels, M.J., Gaylets, C., Kim, N., Lynch, B., Sultan, A., and Mordue, D.G. (2016). The FIKK kinase of Toxoplasma gondii is not essential for the parasite’s lytic cycle. Int J Parasitol 46, 323–332.

Suvorova, E.S., Francia, M., Striepen, B., and White, M.W. (2015). A novel bipartite centrosome coordinates the apicomplexan cell cycle. PLoS Biol 13, e1002093.

Tabb, D.L., Mcdonald, W.H., and Yates, J.R., 3rd (2002). DTASelect and Contrast: tools for assembling and comparing protein identifications from shotgun proteomics. J Proteome Res 1, 21–26.

Teo, G., Liu, G., Zhang, J., Nesvizhskii, A.I., Gingras, A.C., and Choi, H. (2014). SAINTexpress: improvements and additional features in Significance Analysis of INTeractome software. J Proteomics 100, 37–43.

Torres, J.A., Pasquarelli, R.R., Back, P.S., Moon, A.S., and Bradley, P.J. (2021). Identification and Molecular Dissection of IMC32, a Conserved Toxoplasma Inner Membrane Complex Protein That Is Essential for Parasite Replication. mBio 12.

Tosetti, N., Dos Santos Pacheco, N., Bertiaux, E., Maco, B., Bournonville, L., Hamel, V., Guichard, P., and Soldati-Favre, D. (2020). Essential function of the alveolin network in the subpellicular microtubules and conoid assembly in Toxoplasma gondii. Elife 9, e56635.

Treeck, M., Sanders, J.L., Elias, J.E., and Boothroyd, J.C. (2011). The phosphoproteomes of Plasmodium falciparum and Toxoplasma gondii reveal unusual adaptations within and beyond the parasites’ boundaries. Cell Host Microbe 10, 410–419.

Trojan, P., Krauss, N., Choe, H.W., Giessl, A., Pulvermuller, A., and Wolfrum, U. (2008). Centrins in retinal photoreceptor cells: regulators in the connecting cilium. Prog Retin Eye Res 27, 237–259.

Wang, X., Qian, P., Cui, H., Yao, L., and Yuan, J. (2020). A protein palmitoylation cascade regulates microtubule cytoskeleton integrity in Plasmodium. EMBO J 39, e104168.

Warrenfeltz, S., Basenko, E.Y., Crouch, K., Harb, O.S., Kissinger, J.C., Roos, D.S., Shanmugasundram, A., and Silva-Franco, F. (2018). EuPathDB: The Eukaryotic Pathogen Genomics Database Resource. Methods Mol Biol 1757, 69–113.

Wickham, H. (2016). ggplot2: Elegant Graphics for Data Analysis. Springer-Verlag New York.

Youn, J.Y., Dunham, W.H., Hong, S.J., Knight, J.D.R., Bashkurov, M., Chen, G.I., Bagci, H., Rathod, B., Macleod, G., Eng, S.W.M., Angers, S., Morris, Q., Fabian, M., Cote, J.F., and Gingras, A.C. (2018). High-Density Proximity Mapping Reveals the Subcellular Organization of mRNA-Associated Granules and Bodies. Mol Cell 69, 517–532 e511.

