## Supplementary matrrial for "A more complex basal complex: novel components mapped to the *Toxoplasma gondii* cytokinesis machinery portray an expanded hierarchy of its assembly and function"

**Supplementary materials**

**Headers to Supplementary Movies**

**Supplementary Movie S1.** SR-SIM 3D rotation of BCC0-smMyc co-expressing endogenously tagged YFP-MORN1 corresponding with Fig. 3d.

**Supplementary Movies S2 and S3.** Time-lapse microscopy of BCC4-mAID-Myc3 parasites expressing endogenously YFP-tagged MORN1, without (Movie S2) and with IAA (Movie S3). IAA was added at t = - 2 hrs. These movies correspond with Fig. 7d.

**Supplementary Movies S4 and S5.** ExM of BCC4-mAID-3xMyc parasites stably endogenously expressing YFP-MORN1 co-stained with α-tubulin (12G10) antiserum. Movie S4 represents the uninduced wild type, whereas Movie S5 was depleted of BCC4. These movies correspond with Fig. 7e.

**Headers to supplementary tables**

**Table S1. Details on the BioID candidates in the prey-prey heatmap of Fig S1.**

**Supplementary Figures**

**Figure S1**

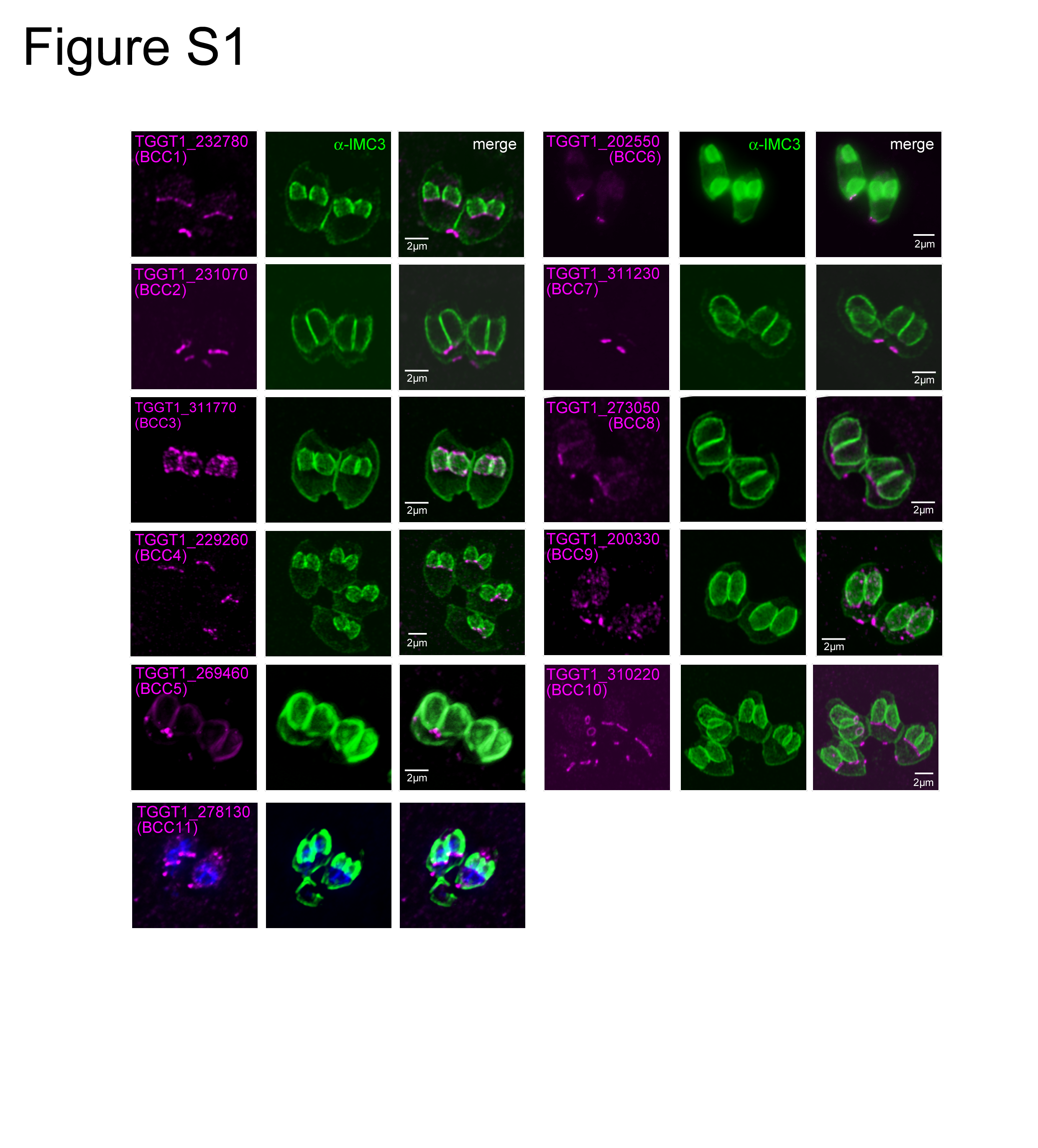

**Fig S1. Localization patterns of BCC components.** All genes were endogenously tagged and co-stained with IMC3 antiserum. All parasite images are of mid-budding to provide a general overview of the spatio-temporal localization dynamics between mother and daughter parasite BC and/or cytoskeleton.

**Figure S2**

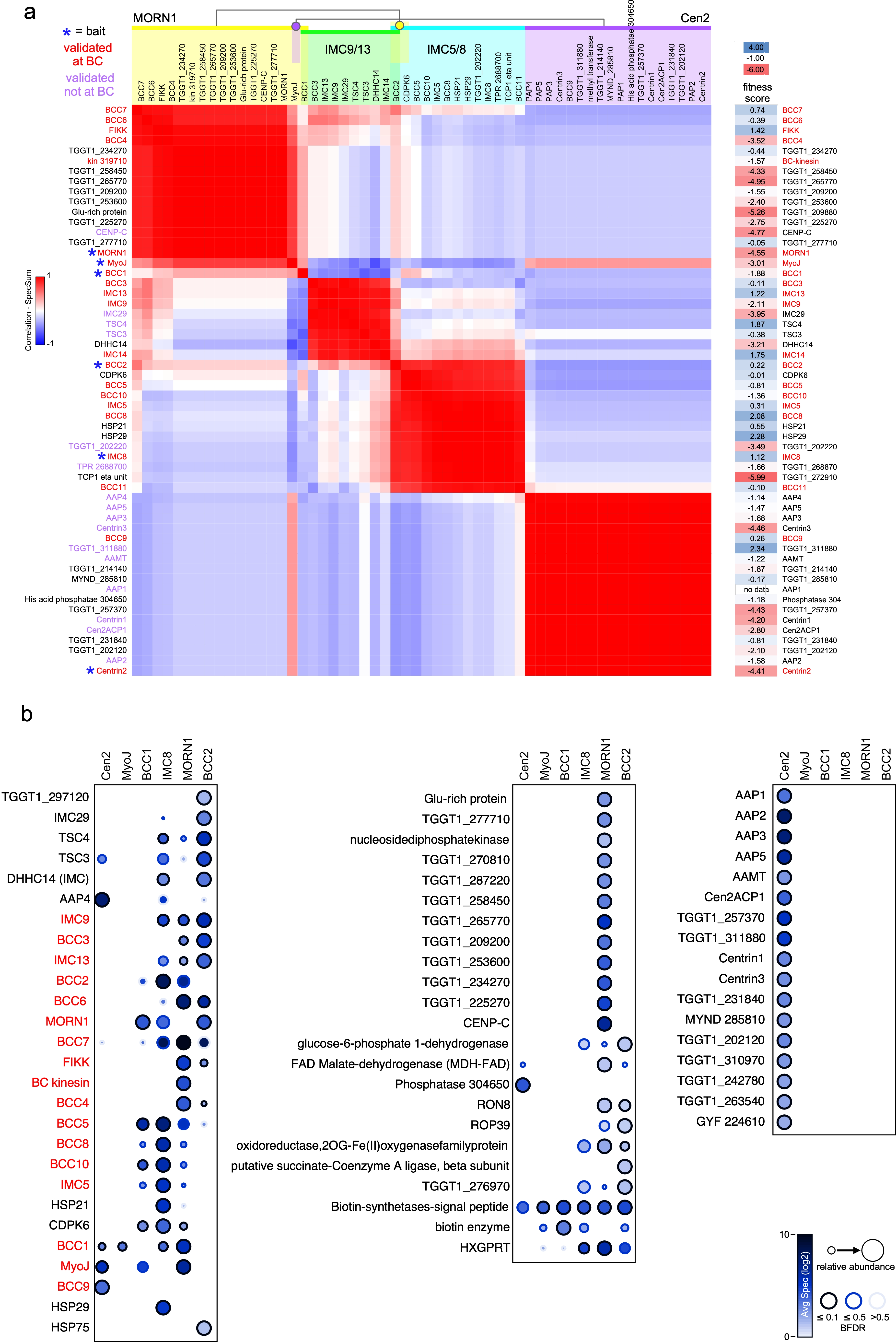

**Fig S2.** **Extended statistical analysis of proximity biotinylation results.**

**a.** Prey-prey correlation map including the BioID2 fusion baits and including cytoplasmic YFP-BioID2 as a negative control. Note that inclusion of this control actually tightens the stringency of proteins assigned to the clusters. BCC0 was discovered by these looser settings shown in Fig. 2a Prohits-viz settings for this correlation map were as follows: Abundance column: “SpecSum”, score column: “False discovery rate (FDR)”, score filter: “0.25”, second score filter: “0.1” and abundance cutoff for prey correlation: “20”. HyperLOPIT, (Barylyuk et al., 2020) and fitness scores (Sidik et al., 2016)for each gene are included on the right side.

**b.** Dotplot including the cytoplasmic YFP-BioID2 as a negative control provides overview regarding the strength of the hits for bait, as indicated. BCC0 and BCC11 did not meet the criteria used to generate this plot. Gene names in red represent experimentally validated BCC components.

**Figure S3**

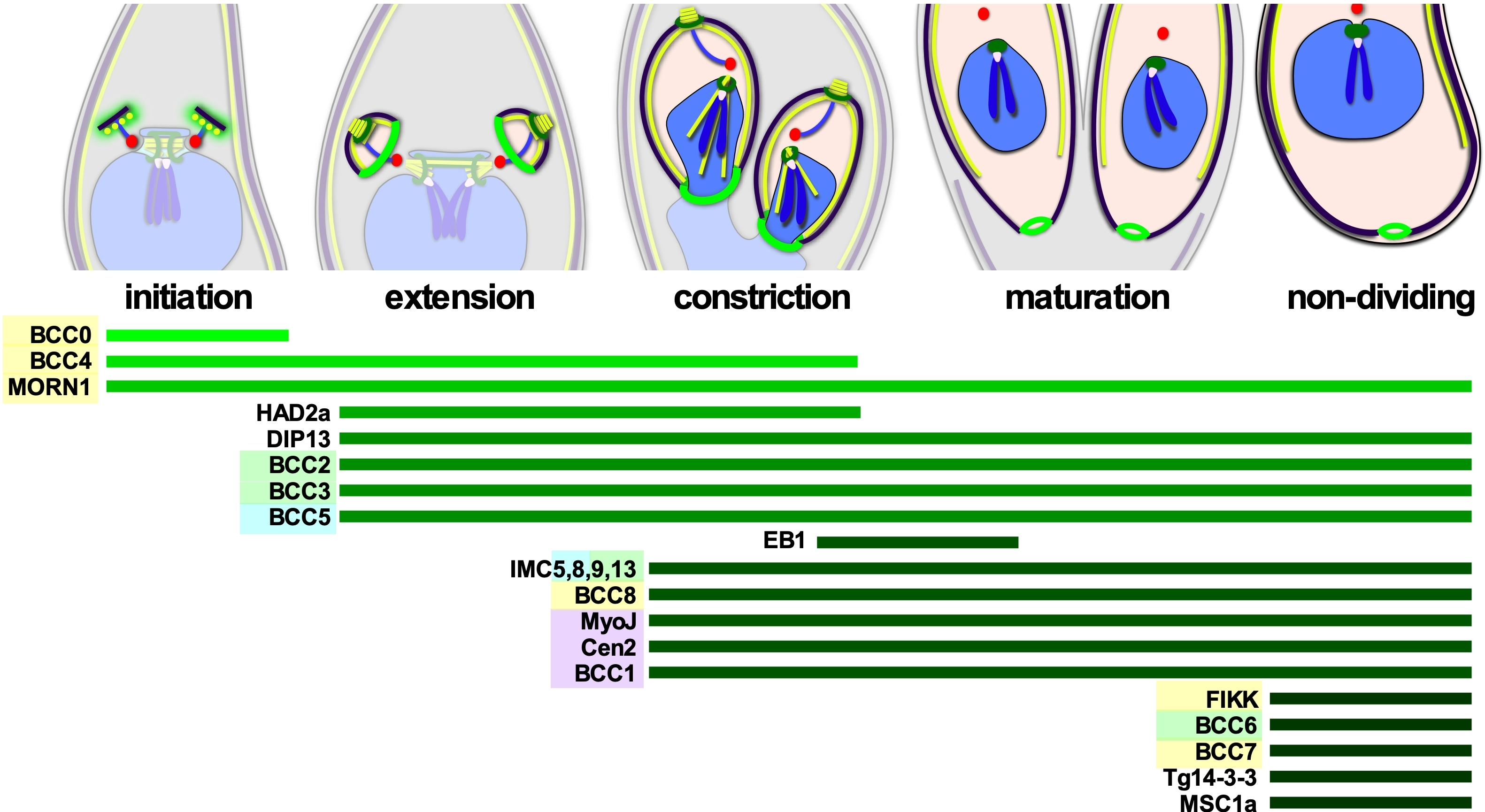

**Fig S3. Temporal patterns of BC composition changes coincide with developing BC function.** Note that BCSCs are largely not assembled sequentially but predominantly in parallel. Colored blocks marking the protein names correspond with the BCSC complexes as follows: yellow, BCSC-1; green, BCSC-2; bleu, BCSC-2; purple BCSC-4. Original data gathered for: HAD2a (Engelberg et al., 2016), DIP13/SSNA1 (Leveque et al., 2016), EB1 (Chen et al., 2015b), IMC5, 8, 9, 13 (Anderson-White et al., 2011), FIKK (Skariah et al., 2016), and finally Tg14-3-3 and MSC1a (Lorestani et al., 2012).

**Figure S4**

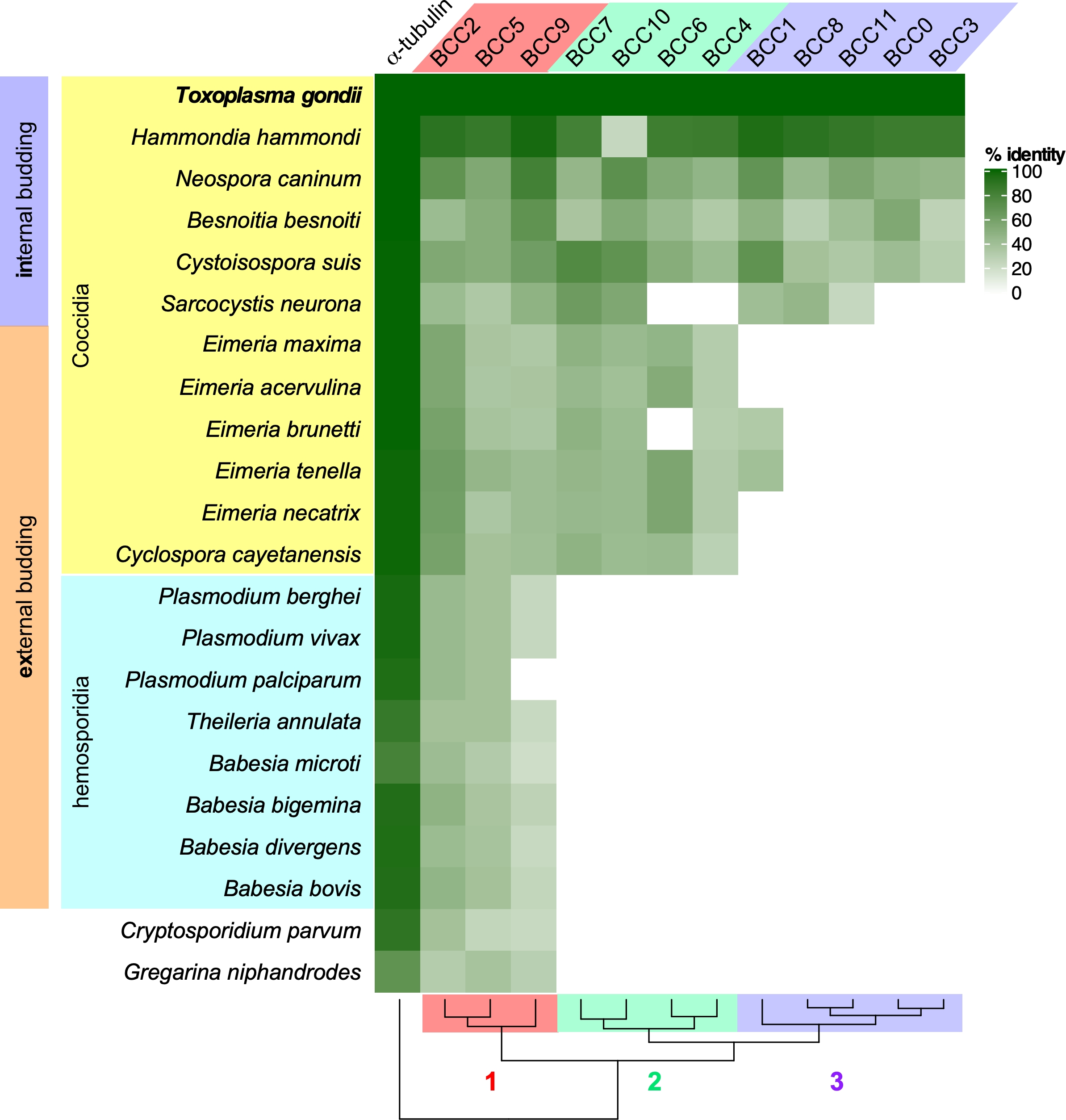

**Fig S4. Gene conservation of newly identified BCCs.** *T. gondii* BCC protein sequences were BLASTP searched for orthologs on EuPathDB (Warrenfeltz et al., 2018) against the following species and strains: *Cryptosporidium parvum* Iowa II, *Cryptosporidium muris* RN66, *Cystoisospora suis* strain Wien I, *Cyclospora cayetanensis* isolate NF1_C8, *Babesia bovis* T2Bo, *Babesia bigemina* strain BOND, *Babesia divergens* strain 1802A, *Babesia microti* strain RI, *Gregarina niphandrodes* Unknown strain, *Theileria annulata* strain Ankara, *Plasmodium falciparum* 3D7, *Plasmodium berghei* ANKA, *Plasmodium vivax* P01, *Toxoplasma gondii* GT1, *Hammondia hammondi* strain H.H.34, *Neospora caninum* Liverpool, *Sarcocystis neurona* SN3, *Eimeria tenella* strain Houghton, *Eimeria maxima* Weybridge, *Eimeria brunetti* Houghton, *Eimeria acervulina* Houghton. Top hits in each species were selected as orthologs. The percentage identify for the orthologs was used to assemble a heatmap

generated in R using the ComplexHeatmap package (Gu et al., 2016). *T. gondii* α-tubulin was used as a reference for a highly conserved gene.
